# Flexible behavioral adjustment to frustrative nonreward in anticipatory behavior, but not in consummatory behavior, requires the dorsal hippocampus

**DOI:** 10.1101/2024.05.22.595420

**Authors:** Christopher Hagen, Megi Hoxha, Saee Chitale, Andre O. White, Pedro Ogallar, Alejandro N. Expósito, Antonio D. R. Agüera, Carmen Torres, Mauricio R. Papini, Marta Sabariego

## Abstract

The hippocampus (HC) is recognized for its pivotal role in memory-related plasticity and facilitating adaptive behavioral responses to reward shifts. However, the nature of its involvement in the response to reward downshifts remains to be determined. To bridge this knowledge gap, we explored the HC’s function through a series of experiments in various tasks involving reward downshifts and using several neural manipulations in rats. In Experiment 1, complete excitotoxic lesions of the HC impaired choice performance in an 8-maze task after reducing the quantity of sugar pellet rewards. In Experiment 2, whereas chemogenetic inhibition of the dorsal HC left consummatory responses unaffected after a sucrose downshift, it significantly disrupted anticipatory behavior following a food-pellet reward reduction. Experiments 3-5 used peripheral lipopolysaccharide (LPS) treatment and found an increase in cytokine levels in the dorsal HC (dHC, Experiment 3), impaired anticipatory choice (Experiment 4), but no effect on consummatory behavior in two reward-downshift tasks. In Experiment 6, after a sucrose downshift, we found no evidence of increased activation in either the dorsal or ventral HC, as measured by c-Fos expression. These findings highlight the HC’s pivotal role in adaptively modulating anticipatory behavior in response to frustrative nonreward, while having no effect on adjustments of consummatory behavior. Spatial orientation, memory update, choice of reward signals of different value, and anticipatory vs. consummatory adjustments to reward downshift are discussed as potential mechanisms that could elucidate the specific effects observed from HC manipulations.

## Introduction

Memories enable organisms to adaptively optimize goal-directed behaviors based on past experience (Biderman, 2020). The hippocampus (HC), recognized as a central hub of memory-related plasticity (Bliss & Gardner-Medwin, 1973; Squire & Kandel, 2009), has been suggested to facilitate adaptive behavioral responses to shifts in rewards within a neural circuit that detects negative disparities between obtained (lower) and expected (higher) rewards (Gray & McNaughton, 2000). In turn, negative mismatches can trigger response inhibition by activating a negative emotional state known as frustrative nonreward (Amsel, 1992). However, the results of experiments involving reward downshifts and HC manipulations reveal inconsistencies.

Franchina and Brown (1971) conducted an instrumental successive negative contrast (iSNC) task wherein rats had to emit an instrumental response to receive a food reward that unexpectedly decreased in value after ten sessions. Extensive HC lesions did not affect acquisition before reward downshift but nullified behavioral changes following the reward devaluation. However, HC lesions failed to uncover any disruption in the consummatory successive negative contrast (cSNC) task, while simultaneously confirming the deficits in the iSNC task (Flaherty et al., 1998; Flaherty et al., 1989; Kramarcy et al., 1973). The inconsistent findings across studies and behavioral paradigms underscore the need to further investigate whether the HC is necessary for animals to adjust to changes in reward conditions.

The fundamental distinction between instrumental and consummatory tasks lies in the animal’s engagement with the reward: iSNC tasks involve an instrumental, anticipatory action driven by reward expectancy, while cSNC tasks center on reward intake. This divergence in task requirements and the corresponding effects of HC lesions on these tasks align with the notion that HC engagement is critical for goal-directed behavior necessitating spatial navigation. HC neurons encode the animal’s current location constructing a map-like representation of the environment (O’Keefe, 1976). Reward delivery then influences these hippocampal maps. For example, cells in the HC cluster their place fields around rewards (Dupret et al., 2010; Poucet & Hok, 2017), and increase firing near reward locations (Poucet & Hok, 2017). Furthermore, some place neurons show systematic changes in firing rate and preferred firing sites across different environments in response to shifts in reward locations (Dupret et al., 2010; Krishnan et al., 2022; Poucet & Hok, 2017). Indeed, neurons in the dorsal HC (dHC) that project to the nucleus accumbens display amplified representations of task-relevant information, encompassing both spatial and non-spatial aspects (Barnstedt, 2023), that play a role in selecting actions and promoting effective appetitive behavior (Barnstedt, 2023; Trouche et al., 2019). Additionally, awake sharp-wave ripple (SWR) events, which coincide with bursts of HC place cell sequences, are enhanced in response to rewards (Singer & Frank, 2009). These reward associated SWRs activate specific place cells linked to reward processing that could contribute to the formation of associations between spatial paths and rewards. Although past studies have primarily emphasized spatial aspects over the appetitive dimension (Sosa & Frank, 2018), these findings collectively suggest a pivotal role of the HC in instrumental goal-directed behaviors that necessitate spatial navigation. However, the specific conditions determining the involvement of the HC in guiding behavioral adjustments to reward downshift remain largely unknown.

The present experiments have two objectives. First, to assess the role of the HC in the cSNC task using different techniques to influence HC circuits. Experiments 2, 4, and 6 were designed to either confirm or rectify the conclusion that the cSNC effect does not depend on HC integrity. Second, to extend the examination of HC function to free-choice situations involving reward downshifts in instrumental behavior in an 8-maze (iSNC, Experiment 1) and in a Pavlovian autoshaping task (pSNC, Experiments 2 and 5). These experiments offer a fresh view of HC function in situations involving negative disparities between obtained and expected rewards that require behavioral flexibility and are associated with frustrative nonreward.

## Experiment 1: Complete HC lesions and instrumental performance in the 8-maze

This experiment explored the effects of complete excitotoxic lesions of the HC on an instrumental task involving a choice between alternatives that differed in terms of reward magnitude. Using an 8-maze task, rats received training in forced-choice trials in which only one option was available, either a right (R) or left (L) turn, each paired with either 12 or 2 pellets (counterbalanced, e.g., R12, L2). After initial acquisition, animals were exposed to free-choice trials in which both options were available (e.g., R12 vs. L2). A preference for the option paired with the largest reward was expected. Eventually, the option paired with the largest reward was downshifted, such that both turns led to the small reward (e.g., R2, L2). Reward downshift was expected to either eliminate the preference for the large-reward option or switch preference to the unshifted option (e.g., either R2∼L2 or R2<L2; Conrad & Papini, 2018). The key question was whether HC lesions would affect free-choice preference after reward downshift.

## Method

### Subjects

The subjects were 14 experimentally naïve, female, Long-Evans rats weighing between 300-350 g, purchased from Charles River Laboratories (Wilmington, MA). Rats were housed individually in ventilated cages with cardboard tunnels and Wood Gnawing Blocks as enrichment devices. Rats were maintained on a reversed 12 h light/dark cycle at 23 ^◦^C. Behavioral testing was performed in the dark phase of the light-dark cycle. During periods of behavioral testing, rats were food restricted and maintained at 85% of their ad libitum weight. The mean (±SEM) ad libitum weight for the entire sample was 327.8 g (±6.1). Water was provided ad libitum in all the experiments included in this article.

Rats were randomly assigned to one of two groups: Group HC (*n*=7) included animals with nearly complete ibotenic acid lesions of the HC and Group Sham (*n*=7) included animals with simulated lesions. All animal experiments were approved by the Mount Holyoke College (Experiment 1), Texas Christian University (Experiments 2-5), and University of Jaén (Experiment 6) Institutional Animal Care and Use Committees and conducted according to National Institutes of Health guidelines.

### Surgery

The general procedure used in Experiment 1 is described in Figure 1a. All surgeries were performed using aseptic procedures. Anesthesia was maintained throughout surgery with isoflurane gas (0.8–2.0% isoflurane delivered in O_2_ at 1 L/min). The animal was positioned in a Kopf stereotaxic instrument and the incisor bar was adjusted until Bregma was level with Lambda. The bone overlying the target site was removed using a high-speed drill. Bilateral excitotoxic hippocampal lesions were produced by local microinjections of ibotenic acid (IBO; Fisher Scientific, Agawam, MA). IBO was dissolved in 0.01 M phosphate-buffered saline (PBS) at a concentration of 10 mg/mL and infused at a rate of approximately 0.1 μL/min using a 10 μL Hamilton syringe (Hamilton, Reno, NV) mounted on a stereotaxic frame and held with a Kopf model 5000 microinjector (David Kopf Instruments, Tujunga, CA). The syringe needle was lowered to the target and left in place for 1 min before the start of the infusion. After the infusion was completed, the syringe needle was left in place for 2 min to reduce the spread of IBO up the needle tract. IBO was injected into 18 sites (total volume 0.51 μL) within each HC (all coordinates are in millimeters and relative to Bregma): anteroposterior (AP) −2.4, mediolateral (ML) ± 1.0, dorsoventral (DV) −3.5; AP −3.2, ML ± 1.4, DV −3.1, −2.3; AP −3.2, ML ± 3.0, DV −2.7; AP −4.0, ML ± 2.5, DV −2.8, −1.8; AP −4.0, ML ± 3.7, DV −2.7; AP −4.8, ML ± 4.9, DV −7.2, −6.4; AP −4.8, ML ± 4.3, DV −7.7, −7.1, −3.5; AP −5.4, ML ± 4.2, DV −4.4, −3.9; AP −5.4, ML ± 5.0, DV −6.6, −5.9, −5.2, −4.5. The procedure for sham animals was the same as for animals with lesions, but the dura was not punctured, the syringe needle was not lowered into the cortex, and IBO was not injected. After completion of each lesion, the wounds were closed, and the animal was allowed to recover from anesthesia on a water-circulating heating pad. Behavioral testing began two weeks after surgery.

**Fig 1.**
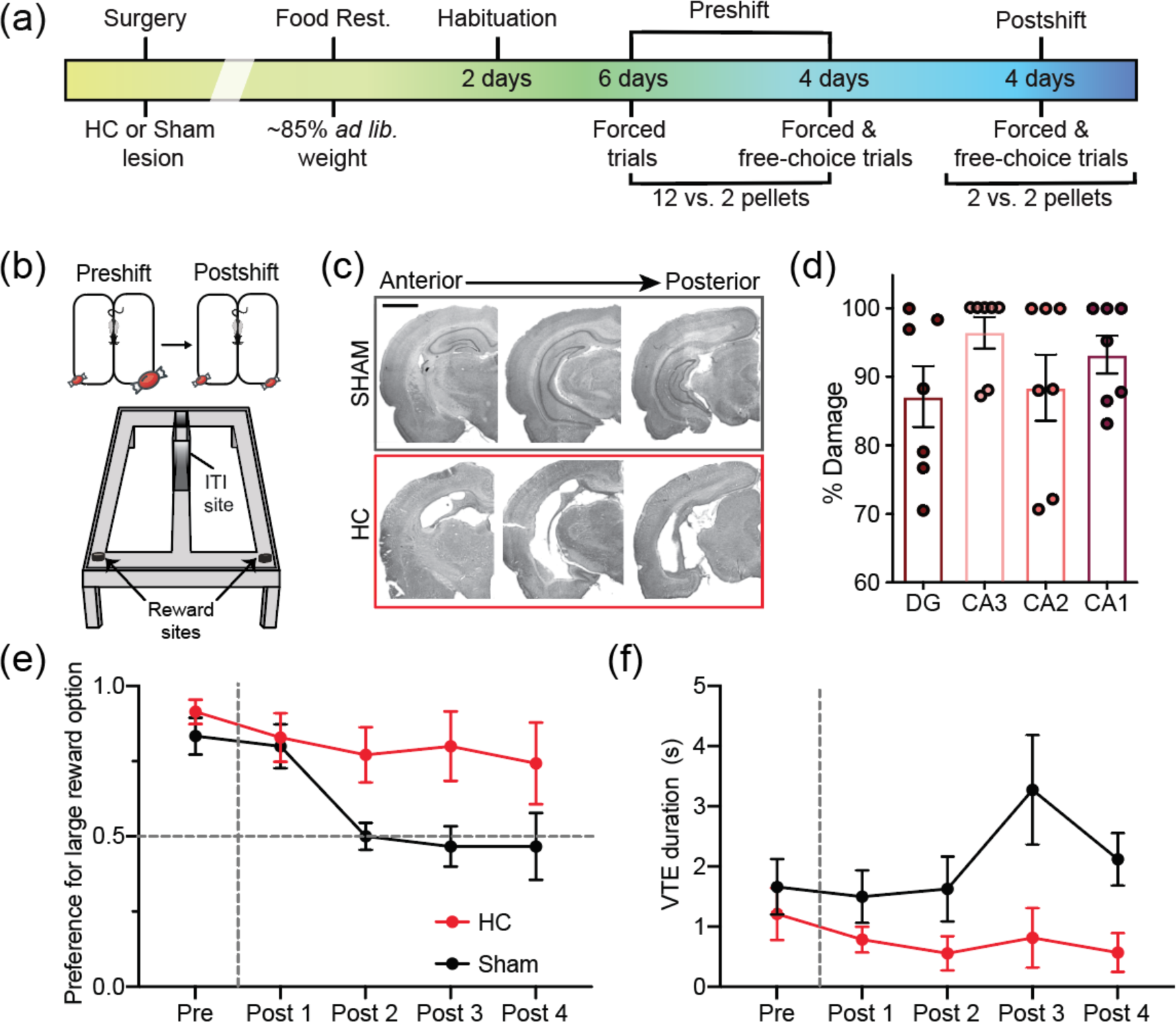
Hippocampal lesions prevent adjustment to reward devaluation in the 8-maze. (a) Experimental timeline. (b) Top: Graphical description of preshift and postshift phases, candy icon size indicates the absolute reward value. Bottom: Illustration of the figure-eight maze with the reward and ITI sites indicated. (c) Photographs through one hemisphere of the rat HC at three coronal levels (anterior to posterior) for rats with sham or HC lesions. Scale bar indicates 2 mm. (d) Average percentage of lesioned tissue, separated by cell layer. (e) Preference for the large reward option in the HC (*n*=7) and Sham (*n*=6) groups. The preference ratio was calculated by dividing the number of visits to the large/devalued reward site by the total number of free choice trials. (f) VTE duration at the choice point. See text for further details.

### Apparatus

Experiment 1 was conducted in a modified T-maze with return arms, giving it an 8-shape design (Figure 1b). The maze was constructed from sanded gray hard polyvinyl chloride plastic runways that were 10 cm wide with 3 cm tall walls on each side of the runway. The center stem of the maze was 120.4 cm long and the cross pieces at each end of the central stem were 90 cm long. The maze was positioned on top of four legs, elevating the base of the maze 68.5 cm off the ground. At one end of the central stem, two movable barriers were positioned 26.3 cm apart serving as the inter-trial interval (ITI) area. The side walls between the two movable doors were 24.5 cm high. There were visual cues around the testing room and the experimenter always stood at the same place in the room relative to the maze. Overhead lighting provided dim illumination in the room.

### Behavior

Behavioral training started after the two weeks of recovery from surgery and involved one session per day. On sessions 1-2 (habituation), rats were handled, were given sugar pellets in their home cages, and were familiarized with the room where the testing would take place by allowing them to freely explore the maze for 10 min with sugar pellets scattered over the floor of the maze. Sessions 3-8 involved preshift training. During preshift training sessions, all rats were exposed to forced-choice trials in which only one direction was available. Thus, turning to the right (R) on right trials led to 12 pellets, whereas turning to the left (L) on left trials led to 2 pellets. The assignment of reward magnitude to each direction was counterbalanced across subjects, but the reward validity of each direction remained constant for any given animal across sessions. Each daily preshift session consisted of 10 trials separated by an ITI of 60 s. On sessions 9-12, rats ran through alternative blocks of five forced-choice trials and five free-choice trials for a total of 10 trials in each session. Reward quantities remained the same during these four sessions, across both forced-choice and free-choice trials, with one site containing 12 pellets and the other containing 2 pellets. Given the odd number of forced trials, each session inherently consisted of a biased set of forced-choice trials, skewing either towards the large or the low reward. We analyzed sessions with varying trial distributions and discovered no significant differences between them. This indicates that the uneven distribution of forced trials did not affect the animals’ preference for the larger reward (Supplementary Figure S1). No correction of choices was allowed during free-choice trials. Free-choice trials were used to assess whether rats had acquired a preference for one of the reward sites. Sessions 13-16 involved postshift sessions. During these sessions, all animals were rewarded with 2 pellets regardless of the arm visited. Each of these postshift sessions consisted of a block of five forced-choice trials followed by a block of five free-choice trials (i.e., 10 trials per session).

### Histology

At the completion of testing, rats were administered an 80 mg/kg of sodium pentobarbital and perfused transcardially with 0.01 M PBS followed by 4% paraformaldehyde solution (in 0.01-M PBS). Brains were then removed from the skull and kept in a solution of 4% paraformaldehyde for 24 h before they were transferred to a 30% sucrose solution where they remained for at least 48 h. Coronal sections (40-μm thick) were cut with a freezing microtome beginning at the level of the anterior commissure and continuing posterior through the length of the HC. Every fifth section was mounted and stained with cresyl violet to assess the size of the lesions. Each section was assessed under magnification using the Cavalieri method on ImageJ. The tissue was considered damaged if it was absent or necrotic (i.e., hippocampal tissue was present, but there was no evidence of Nissl staining, or the tissue was gliotic). The volume of the spared tissue in the dentate gyrus (DG), CA1, CA2, and CA3 cell layers was quantified. The percentage of damage was calculated by normalizing the volume of spared tissue in each animal in Group HC to the average volume of the HC in animals from Group Sham according to the following formula: 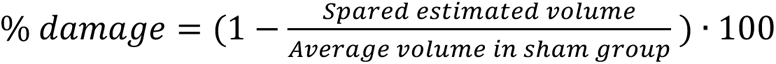

### Statistics

The dependent variables were reward choice (large vs. small, downshifted vs. unshifted), measured by direct observation and by video recording to assess reliability. Vicarious trial and error (VTE) duration was defined as the time at the choice point when the animal is moving its head back and forth over potential paths (right and left alleys). VTE was measured using the ethogram software BORIS (Friar & Gamba, 2016). Videos were coded by two experimenters who were blind to the experimental conditions and the results from their scoring were averaged for statistical analysis and plotting purposes. Video coders received previous training to minimize variability in coding procedures.

In all the statistics reported in this article, Group was always a between-subject factor whereas Session was always a within-subject factor. An alpha value equal or lower than 0.05 was used for statistical inferences, effect size was indexed with the partial eta squared (*η_p_*^2^) statistic, and Bonferroni pairwise comparisons derived from the main analysis were computed to identify the source of significant interactions. To simplify statistical reports, *p* and *η_p_*^2^ values are reported only when *F*>1. All statistics were computed with the IBM SPSS Version 26 package.

## Results

One rat from the Sham group did not consume the reward pellets during preshift sessions and was therefore excluded from the experiment, leaving this group with 6 animals. Representative brain slices show the brains of Sham and HC animals (Figure 1c). The percentage of HC tissue damaged in the DG and CA fields provides an accurate picture of the extent of the lesions (Figure 1d). These lesions were extensive, averaging at least 85% of damage in each of the areas.

Preference for the option paired with the large reward in free-choice trials, for the last preshift session and the four postshift sessions is shown in Figure 1e. Preference was high in preshift, but as expected, reward downshift reduced it in Sham animals. However, rats with HC lesions persevered in their preshift preference. A Group by Session analysis revealed a significant interaction, *F*(4, 44)=2.662, *p*=0.045, *η_p_*^2^=0.195, and a significant change across sessions, *F*(4, 44)=8.868, *p*<0.001, *η_p_*^2^=0.446, but the group effect was nonsignificant, *F*(1, 11)=3.651, *p*=0.082, *η_p_*^2^=0.249. Pairwise Bonferroni tests indicated that the source of the significant interaction was a significantly higher preference for the large-reward option in HC animals over Sham animals on the second and third postshift sessions, *p*s<0.04.

A similar difference was observed when preference was assessed only during the first free-choice trial of each session, when carry-over stimuli from previous trials is unlikely to play a significant role given the 24-h interval between sessions. The mean preference for the large reward options during the last two preshift sessions was 0.83 for Shams and 0.92 for HC animals. However, whereas this preference decreased during the last two postshift sessions to 0.46 for Shams, it remained high for HC rats at 0.77. Thus, after two sessions of reward downshift, the choice of the large-reward side was reduced in Shams, while HC animals remained committed to the response preference acquired during preshift sessions.

The HC lesion also affected the duration of VTE behavior (Figure 1f). Sham animals showed consistently higher levels of VTE than HC animals, especially during postshift sessions. These results yielded a significant Lesion effect, *F*(1, 11)=17.53, *p*=0.002, *η_p_*^2^=0.61, although the Session and Lesion by Session interaction were nonsignificant, *F*s<1.94, *p*s>0.19, *η_p_^2^*s<0.15. To further explore VTE difference between groups, we conducted additional analyses focusing on individual trial types. Independent-sample t-tests comparing VTE duration between the HC lesion and Sham groups on the last day of preshift showed no significant difference, *t*(11)=0.71, *p*=0.49, indicating that the lesion effect was not evident during preshift trials. Conversely, a significant difference was observed on postshift session 3, *t*(10)=2.56, *p*=0.0285, underscoring a specific lesion effect during this phase.

HC lesions reduced the level of VTE behavior during postshift free-choice trials, but did not affect the preference for the option paired with the large reward during preshift trials. This reduction in VTE behavior is in agreement with previous studies indicating the dorsal HC’s crucial role in the deliberation process of decision-making (Meyer-Mueller et al., 2020) and the modulatory effect of clonidine on hippocampal choice processing to enhance decisiveness (Amemiya & Redish, 2016). Importantly, HC lesions led to significant perseverance of the initial preference after reward downshift was introduced. This result is reminiscent of the effects of HC lesions on appetitive extinction (e.g., Jarrard, Isaacson, & Wickelgren, 1964; Micco, McEwen, & Shein, 1979) and in progressive ratio schedules (Schmelzeis & Mittleman, 1996). In these early studies, HC lesions increased response perseverance during extinction in a runway task.

## Experiment 2: Chemogenetic inhibition of dHC and behavior in the consummatory reward downshift (cRD) and pSNC tasks

The massive HC lesions used in Experiment 1 extended previous research on reward downshift in runway situations (Franchina & Brown, 1971) to a free-choice task. In these two cases, HC lesions rendered the behavior resistant to change after the downshift, without affecting the ability to acquire the behavior during preshift sessions. Experiment 2 sought to extend these results using a chemogenetic tool known as designer receptors exclusively activated by designer drugs (DREADDs). These are engineered muscarinic receptors modified to respond to an exogenous ligand (e.g., clozapine-N-oxide, CNO) injected systemically before specific training sessions. In this experiment, an inhibitory DREADD [hM4D(Gi)] was infused in the dorsal hippocampus (dHC), thus limiting the location of the lesion relative to the more complete lesions of the previous experiment. In addition, animals were exposed to two tasks involving reward downshift in successive phases. During Phase 1, rats received access to 32% sucrose for 10 5-min sessions and were subsequently downshifted to 2% sucrose for 3 additional sessions, while their consummatory behavior (licking at a sipper tube delivering sucrose) was recorded (Flaherty, 1996). Because unshifted controls were not included in Experiments 2 and 4, we refer to the behavioral effect as cRD, rather than cSNC. The cSNC effect is defined as a deterioration of behavior beyond the level of an unshifted control (Flaherty, 1996), whereas the cRD effect emphasizes the suppression of behavior that follows the unexpected reward downshift. Either CNO or vehicle (Veh) injections were administered before postshift sessions 11 and 12. The primary purpose of the cRD task used in this experiment was to study the effect of chemogenetic inhibition of the dHC on the suppression of consummatory behavior that follows reward downshift. Based on previous lesion results (e.g., Flaherty et al., 1989; Kramarcy et al., 1973), chemogenetic inhibition should produce little or no effect on consummatory behavior after a reward downshift.

In Phase 2, all the rats were exposed to a Pavlovian autoshaping task (pSNC) in which two levers located at either side of the food container were paired with either 12 or 2 pellets (e.g., R12, L2; counterbalanced across subjects) in sessions involving 3 forced-choice trials with each lever. Some of these sessions ended with a nonreinforced free-choice trial with simultaneous access to both levers (e.g., R vs. L). It was expected that rats would respond more to the lever previously paired with the large reward than to the lever paired with the small reward (i.e., R>L in this example). In a final (postshift) session, one of the levers was downshifted to the small reward (e.g., R12-2), while the other remained unshifted (e.g., L2-2). The effects of this reward downshift were assessed in a nonreinforced free-choice trial equal to that administered at the end of preshift training (i.e., R vs. L). The expected effects of such downshift were to either reverse the lever preference (R<L; Conrad & Papini, 2018) or to eliminate the initial preference (R∼L). To the extent that pSNC and iSNC (Experiment 1) tasks are based on anticipatory behavior, it was expected that chemogenetic inhibition would induce perseverance of preference for the lever paired with the large reward after the downshift.

## Method

### Subjects

The subjects were 28 male Wistar rats, approximately 90 days old at the start of the experiment, experimentally naïve, born from breeders purchased from Charles River Laboratories (Wilmington, MA) and reared under standard conditions at the Texas Christian University (TCU) Vivarium Research Facility. Pups were weaned at about 21 days of age and transferred to single-sex polycarbonate cages in groups of at least two animals. At around 40 days of age, they were transferred to wire-bottom cages, each containing a rodent retreat for enrichment, where they remained until the end of the experiment. Animals were fed with standard laboratory rat chow, freely available until reaching 90 days of age, when food restriction was implemented. These developmental conditions were common to all animals used in the experiments conducted at TCU. Ad libitum weights (g) for individual animals were calculated by averaging the weight of the animal across two consecutive days immediately prior to food restriction. The mean (±SEM) ad libitum weight for the selected animals was 483.5 g (±7.2). In preparation for surgery, access to food was restricted until rats reached about 90% of their ad libitum weight. This initial food restriction was used to reduce the number of days before the start of behavioral testing (see Kawasaki et al., 2017). In preparation for behavioral testing, weights were further reduced to an 81-84% of the ad libitum weight for each animal. All animals received food every day, but the amount was determined individually to keep weights constant across the experiment. During the experiment, animals were under a 12 h light/dark cycle (lights on at 07:00 h), in a colony room with constant temperature (22-23°C) and humidity (45-65%).

### Surgery

The general procedure of Experiment 2 is described in Figure 2a. To induce anesthesia, the animals were placed in a chamber filled with a mixture of breathing air and isoflurane vapor (5% for induction, 1-2% for maintenance). When their breathing grew deep and slow, the animal was positioned in a stereotaxic frame (Angle Two, Leica, program version 3.0.0; Leica Biosystems, Buffalo Grove, IL) that maintained the delivery of isoflurane vapor to keep the animal under anesthesia during surgery. The frame was fixed with blunt-tipped ear bars, a bite bar, and a mask. Once the animal was situated in the frame, its eyes were covered with Vaseline to prevent eye dryness and possible harm from the microscope light. The area of the scalp to be incised was shaved and glazed with Betadine (povidone-iodine topical solution, 10%), after which an incision was made at midline of the scalp. Blunted hooks were used to separate the two sides of the incision and bare the skull, which was then carefully cleaned. Protective layers were peeled back from the surface. After assessing and adjusting the position of the skull for flatness, the location of the dHC was determined based on the Paxinos and Watson (2013) atlas and marked on both sides of the skull. Each marked site was drilled and infused with the viral construct using a 10-μl Hamilton syringe fixed on a stereotaxic injector (Quintessential Stereotaxic Injector, Stoelting, Wood Dale, IL), administering 0.5 μL of virus per site at a rate of 0.15 mL/min. There were two infusion sites per hemisphere for a total of four infusions at HC coordinates (AP -3.1, ML ±0.7, DV -3.7, and AP -3.6, ML ±0.75, DV -4.0). The Hamilton syringe remained in place for an additional 10 min for the fluid to diffuse into the brain tissue, after which the syringe was slowly withdrawn and the scalp was stapled back together to facilitate healing. The animal was then removed from the stereotaxic frame and injected with buprenorphine hydrochloride (0.05 mg/kg, subcutaneous) to dampen surgery-induced pain; a second dose was administered 24 h after surgery. During the ensuing 5-day recovery period, animals stayed individually housed in polycarbonate cages, where they had access to food at a 90% restriction level, including typical lab rodent chow and supplementary recovery gel, as well as readily available water. At the end of the recovery period, animals were returned to their typical wire-bottom home cages and food restricted to an 81-84% of their ad libitum weight in preparation for behavioral testing.

**Fig 2.**
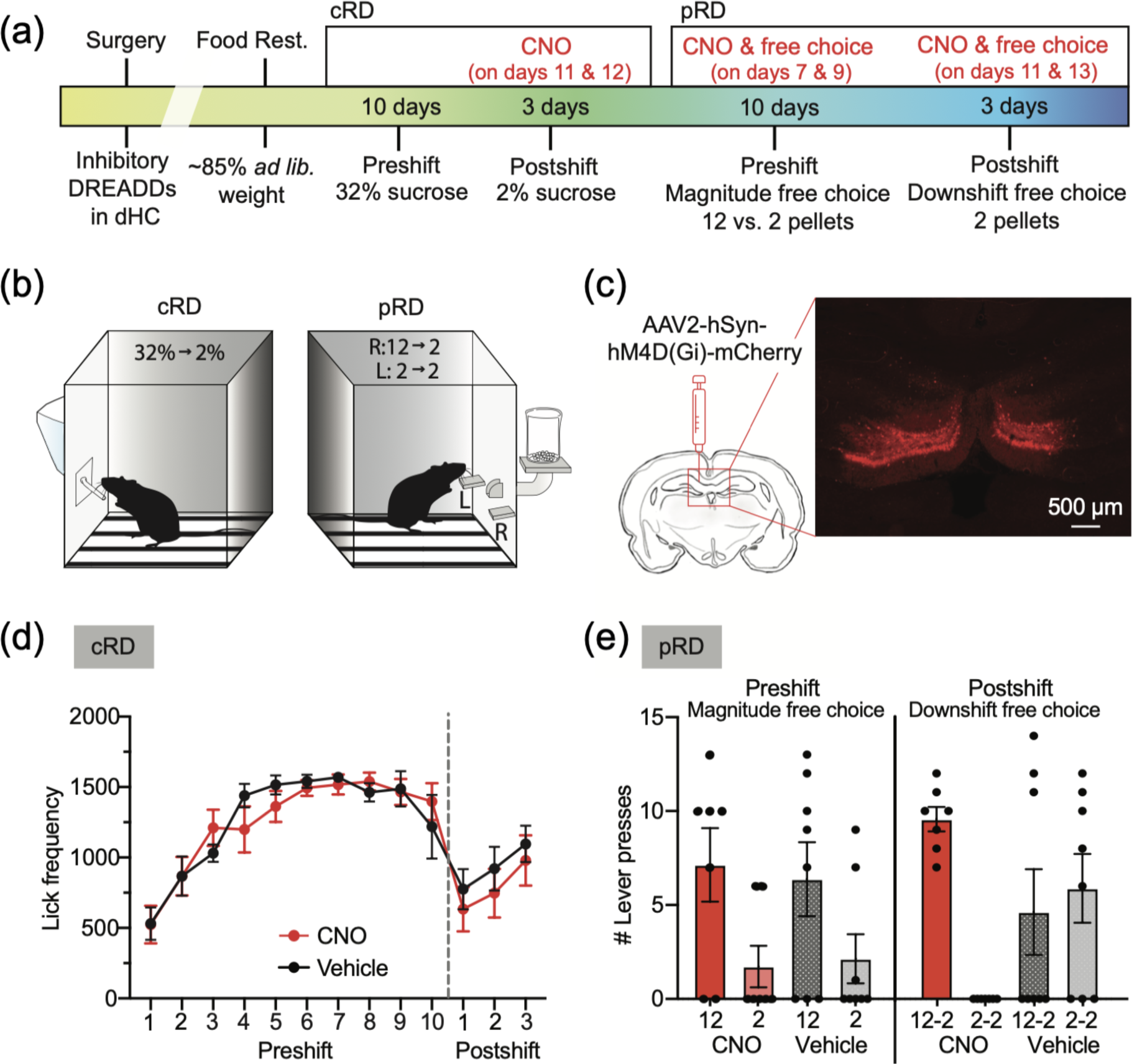
Chemogenetic inactivation of the dorsal HC prevents adjustment to reward devaluation in anticipatory behavior, but not in consummatory behavior. (a) Experimental timeline. (b) Illustration of the behavioral apparatus for the cRD task (left) and the pSNC (right). (c) Illustration indicating the injection site and representative brain section photograph showing dorsal hippocampal neurons transfected with the DREADD virus AAV2-hSyn-hM4D(Gi)-mCherry. Scale bar indicates 500 μm. (d) Average lick frequency on each experimental day for CNO (*n*=7) and vehicle (*n*=8) groups. (e) Average number of lever presses to each lever per group during preshift (left) and postshift (right). See text for further details.

### Viral vector constructs

The inhibitory [hM4D(Gi)] DREADDs were delivered into the dHC via intracranial infusions of an adeno-associated virus AAV2. The viral vector constructs (inhibitory: pAAV-hSyn-hM4D(Gi)-mCherry 3×10¹² virus molecules/mL, catalog # 50474-AAV2; Addgene, Cambridge, MA) contain a red fluorescent reporter (mCherry) and a DNA fragment for an engineered muscarinic receptor, M4, that responds selectively to the activator drug for DREADDs. The activator drug, CNO, binds to the designer receptor and inhibits neural activity.

Two control conditions that failed to influence consummatory behavior under the same conditions used here (see below) were included in other experiments. First, a viral vector control construct (pAAV-hSyn-eGFP, 7×10¹² virus molecule/mL, catalog # 50465-AAV2; Addgene, Cambridge, MA) infused in the nucleus accumbens (Guarino et al., 2023) and receiving either CNO or Veh injections before exposure to a 32-to-2% sucrose downshift. Second, intact animals receiving CNO or Veh injections during exposure to a 32-to-2% sucrose downshift. Thus, neither the virus nor CNO alone affected performance during reward downshift under the behavioral conditions used in this experiment. Because these manipulations had no influence on consummatory behavior they were not included in the present experiment.

### CNO preparation and injection procedure

CNO (3 mg/kg, ip; NIDA Drug Supply Program) was dissolved in 5% dimethyl sulfoxide (DMSO) and 95% sterile saline. CNO and Veh injections were both administered using the same volume (1 mL/kg) and content, except CNO was not included in Veh preparation. Injections were administered 30 min prior to behavioral testing in a room different from that where the training was administered.

### Phase 1: cRD task. Apparatus and procedure

The conditioning boxes used in the cRD and pSNC tasks (Figure 2b) were located in different rooms and were controlled by separate computers located in yet a third room. These computers controlled the presentation of the bottle and recorded licking responses in the cRD task; and controlled the presentation of levers, the delivery of pellets, and the recording of lever presses and magazine entries in the pSNC task. MED Notation software (MED Associates, St. Albans, VT) was used to program training sessions for both tasks.

The cRD task was conducted in eight conditioning boxes (MED Associates, St. Albans, VT) made of aluminum and Plexiglass (29.3×21.3×26.8 cm, L×H×W). Each chamber was placed in a sound-attenuating enclosure equipped with a speaker generating white noise and a fan, both of which produced noise with an intensity of 80.1 dB (SPL, scale C). Diffuse light was provided by a bulb (GE 1820) located in the center of each enclosure’s rear wall. The floor was made of steel rods running parallel to the feeder wall. A tray with corncob bedding was placed below the floor to collect feces and urine. A hole 1-cm wide, 2-cm long, and 4 cm from the floor was located in the feeder wall; a sipper tube (1 cm in diameter) was automatically inserted through this hole. When fully inserted, the sipper tube was about 2-3 mm outside the internal wall, so animals had to extend their tongue to reach the tip of the sipper. When a rat contacted the sipper tube, a circuit involving the steel rods in the floor was closed and lick frequency was automatically recorded. Lick frequency was defined as the total number of licks recorded during the 5-min session. A computer located in an adjacent room controlled the presentation and retraction of the sipper tube, and recorded lick frequency.

cRD training started once animals reached the target weight range and at least 10 days following viral infusion to optimize DREADD expression. Animals were randomly assigned to one of two groups, both infused with inhibitory DREADDs: CNO vs. Veh injections, administered before each of the three postshift sessions. For all the animals, preshift sessions 1-10 involved access to 32% sucrose whereas postshift sessions 11-13 involved access to 2% sucrose. Animals received training 7 days per week, at about the same time every day (between 10:00 and 13:00 h), during the light portion of the daily light/dark cycle, at a rate of one session per day. Sessions lasted 5 min from the first detection of a sipper-tube contact. CNO or Veh injections were administered in a separate room 30 min before postshift sessions 11-12. Sucrose solutions (32% and 2%) were prepared every 2-3 days, weight/weight by mixing 68 g (or 98 g) of deionized water for every 32 g (or 2 g) of commercial extra-fine granulated pure cane sugar (Imperial Sugar, Imperial Sugar Company, Sugar Land, TX). The mixture containers were shaken until the sugar was completely dissolved. Sucrose solutions were kept at room temperature between sessions.

### Phase 2: pSNC task. Apparatus and procedure

The pSNC task (autoshaping procedure) was conducted in four standard operant chambers (MED Associates, St. Albans, VT). Each chamber measured 20.1×28×20.5 cm (L×H×W) and was contained in a sound-attenuating enclosure equipped with a house light (GE 1820), a speaker to emit white noise, and a fan to promote airflow. The speaker and fan combined produced background noise with an intensity of 80.1 dB (SPL, scale C, Digital Sound Lever Meter, Extech, Waltham, MA). The floor consisted of 0.4-cm diameter stainless steel bars spaced 1.6 cm apart (from center to center). A tray filled with corncob bedding was placed underneath the bars to collect droppings, which were removed as necessary after each session. Precision pellets (45 mg, Bio-Serv, Flemington, NJ) were delivered from an external pellet dispenser into a food cup placed inside a small hole located in the center of the front wall, 2 cm above the floor. Food cups were equipped with photocells located 1.1 cm inside the hole and above the food cup. These photocells detected head insertions into the hole (magazine entries). Located 1 cm at either side of the food cup and 6 cm above the floor were two stainless steel retractable levers. Levers were 4.8 cm wide and when fully inserted protruded 1.9 cm into the chamber. It took 0.2 s for the lever to be fully inserted or retracted. Levers were adjusted so that they could be pressed with a minimum force (lever presses).

Training in the pSNC task started the day after the last session in the cRD task and lasted 13 sessions, one session per day, 7 days per week. In preshift sessions 1-10, the presentation of one lever for 10 s signaled the free delivery of 12 food pellets and the presentation of the other lever for 10 s signaled the free delivery of 2 pellets. Because autoshaping is a Pavlovian procedure, lever pressing (or any other response) was not necessary to produce food delivery. Nevertheless, rats tend to approach and touch, bite, lick, and direct several responses to the lever that result in movements that are recorded as lever presses (i.e., sign tracking; Hearst & Jenkins, 1974). Rats also insert their heads into the hole where food is to be dispensed, a behavior recorded as magazine entries (i.e., goal tracking; Boakes, 1977).

Each preshift session (sessions 1-10) involved 6 forced-choice trials, including 3 right-lever and 3-left lever trials, presented in a random sequence. On each forced-choice trial, one of two levers was presented for 10 s and its retraction coincided with the delivery of food. Pairing of reward magnitude to levers was counterbalanced. For some animals, the right lever was paired with 12 pellets and the left lever was paired with 2 pellets, whereas for others, the lever-pellet pairings were the opposite. Pellets were delivered automatically at the end of each trial, when the lever was retracted, at a rate of one pellet every 0.2 s. On sessions 7 and 9, in addition to the 6 forced-choice trials described above, there was a single free-choice trial in which both levers were presented simultaneously for 10 s and lever presses to each of them were recorded. No food was delivered at the end of these free-choice trials. On postshift sessions 11-13, the 12-pellet lever was downshifted to 2 pellets (downshifted lever), while the 2-pellet lever stayed constant (unshifted lever). The downshift sessions also included 6 forced-choice trials, 3 with each lever, but now every trial ended with the delivery of 2 pellets. In addition, sessions 11 and 13 included a single nonreinforced free-choice trial with both levers available simultaneously for 10 s presented at the end of these downshift sessions. In both forced-choice and free-choice trials, animals could respond to the available lever(s) or not respond at all. CNO or Veh injections were administered in a separate room 30 min before preshift sessions 7 and 9, and also before postshift sessions 11 and 13. We used a predetermined number of sessions for postshift free-choice trials in the present pSNC experiments (Experiments 2 and 5) because the pretrial manipulations (DREADDs and LPS) could not be repeated for several sessions until each animal showed the expected reversal of preference. Under these conditions, it was expected that not all animals would show clear preference for the unshifted lever after the downshift as this tends to develop across free-choice trials (Conrad & Papini, 2018). Instead, we expected a mixture of preferences across animals after the postshift, some still preferring the lever formerly paired with 12 pellets (now downshifted to 2 pellets) and some showing a preference for the unshifted lever.

At the end of each session, in both phases, animals were placed back in their home cage and transported to the colony room. A wet sponge was used to wipe the conditioning boxes after each session. Feces were removed and bedding replaced as needed.

### Histology

After the last pSNC session, rats were anesthetized with sodium pentobarbital and transcardially perfused with 0.1 M PBS followed by 4% paraformaldehyde in 0.1-M PBS, pH 7.4. Immediately after extraction, brains were placed in 4% paraformaldehyde for at least 3 days and then placed in 30% sucrose for at least 2 days for cryoprotection. Brains were sectioned coronally in 40-μm thick slices using a cryostat (Leica Biosystems, Buffalo Grove, IL) and placed directly onto slides. Fluoromount-G mounting medium and cover slips were applied on the slides to preserve the mCherry fluorescent tag. The location of the virus was assessed via fluorescence microscopy (Nikon eclipse Ti inverted microscope, Nikon, Melville, NY) and images were processed using CRi Nuance FX multispectral imaging system (Caliper Life Sciences, Hopkinton, MA) and Nuance 3.0 imaging software (Caliper Life Sciences, Hopkinton, MA).

### Statistics

Lick frequency, the dependent variable in the cSNC task, was subjected to conventional analysis of variance, as described in Experiment 1. Bayesian repeated measures analyses of variance were also computed using JASP Version 0.18.3 to aid in the interpretation of null findings. Bayes Factors (BFs) were estimated for the Group and Group-by-Session interaction effects by comparing models that included those effects with models that only included Session. BFs quantify the evidence in favor of an effect. For example, a BF of 10 for Group would indicate that the data are 10 times more likely to support a model that hypothesizes the presence of a group effect compared to a Session-only model, whereas a BF of 0.10 for Group (i.e., 1/10) would indicate that the data are 10 times more likely under a Session-only model that hypothesizes the absence of a group effect. A BF of 1 would indicate that data are equally likely under either hypothesis, which suggests results are inconclusive. Thus, in the case of null findings, BFs can be used to distinguish between evidence of no group difference and inconclusive evidence. The range of 0.33<BF<3 is commonly considered inconclusive (Keysers et al., 2020).

Two dependent variables were recorded in the pSNC task: lever presses per trial and magazine entries per trial. In turn, these variables were recorded separately for forced-choice and free-choice trials. Given that their frequency is not homogeneous (see Conrad & Papini, 2018), lever presses in forced-choice trials occur at relatively high frequency and were thus subject to conventional analysis of variance as described previously. However, for lever presses in free-choice trials and magazine entries in both forced- and free-choice trials, the frequencies can be very low with many zeroes, thus violating the equal variance assumption. Consequently, they were analyzed using the Wilcoxon signed ranks nonparametric tests for paired samples. All Wilcoxon tests reported in this article were two-tailed. The alpha value was still kept at 0.05.

## Results

### Histology

Based on histological evidence, 13 rats were discarded for lack of clear fluorescence localization, absence of fluorescence in one hemisphere, or fluorescence present outside the dHC. The remaining 15 animals were distributed as follows: Group CNO *n*=7 and Group Veh *n*=8. Figure 2c shows a slice of one of the selected animals mapped to an atlas image. Supplementary Figure S2 shows additional images. DREADDs were localized mainly in the DG.

### cRD task

As expected in the cRD task, licking frequency increased across preshift sessions 1-10 to reach a relatively high and stable level, followed on session 11 by a drastic reduction of licking, which subsequently showed some degree of recovery (Figure 2d). An analysis of these results failed to reveal a significant Group effect (CNO vs. Veh) or a significant Group by Session interaction, either during preshift (Group BF=0.34, Group by Session BF=0.05) or postshift session, *F*s<1 (Group BF=0.53, Group by Session BF=0.14). There was a significant increase in licking across preshift sessions, *F*(9, 117)=22.50, *p*<0.001, *η_p_^2^*=0.63, but the increase in postshift recovery was only marginal, *F*(2, 26)=3.13, *p*=0.061, *η_p_^2^*=0.19. Thus, there was evidence of acquisition of licking during preshift sessions, but only a trend toward recovery of licking after an initial suppression following sucrose downshift. An additional analysis was conducted comparing sessions 10 (last preshift) vs. 11 (first postshift) to determine whether the 32-to-2% sucrose downshift affected behavior. There was a significant degree of consummatory suppression from session 10 to 11, *F*(1, 13)=10.29, *p*<0.008, *η_p_^2^*=0.44, but the Group and Group by Session interaction effects were nonsignificant, *F*s<1 (Group BF=0.41, Group by Session BF=0.24).

### pSNC task

As expected based on previous results (Conrad & Papini, 2018), forced-choice trials yielded no evidence of differential lever pressing performance, whether in preshift or postshift. A Phase (preshift, postshift) by Lever (12, 2) by Group (CNO, Veh) analysis detected nonsignificant differences for all factors, *F*s<1.44, *p*s>0.25, *η_p_^2^*s<0.10.

For free-choice trials, there were a large number of zero responses as animals typically responded to only one of the two levers. During preshift free-choice trials (reward magnitude), CNO animals showed a marginal preference for the 12-pellet lever, *Z*=-1.88, *p*=0.061, whereas Veh animals showed a nonsignificant preference trend, *Z*=-1.404, *p*=0.16. During postshift free-choice trials (reward downshift), CNO animals exhibited a significant preference for the downshifted lever over the unshifted lever, *Z*=-2.371, *p*=0.018, but Veh animals exhibited no preference, *Z*=-0.140, *p*=0.888. If only animals that passed the magnitude test are considered, that is, animals that responded more to the 12-pellet lever than to the 2-pellet lever in the free-choice test (Figure 2e), then CNO animals exhibited a preference for the 12-pellet lever over the 2-pellet lever in preshift, *Z*=-2.060, *p*=0.039, and that preference was maintained after the 12-to-2 pellet downshift, *Z*=-2.023, *p*=0.043. By contrast, Veh animals showed a significant preference for the 12-pellet lever in preshift, *Z*=-2.023, *p*=0.043, but that preference was eliminated in postshift, *Z*=-0.405, *p*=0.686.

In our experiments using the pSNC procedure, we typically observe an early increase in goal tracking during the initial 1-4 sessions followed by a sharp decrease in subsequent sessions maintained even through sessions of extinction (e.g., Glueck et al., 2018; Torres et al., 2016). This pattern was also observed in the present experiment. Thus, by the time we administered free-choice trials, magazine entries were already extremely low with a substantial number of zeros in all the trial types. Wilcoxon nonparametric tests for related samples comparing preshift and postshift forced-choice trials, for CNO and Veh conditions, yielded uniformly negative results, *Z*s<-1.360, *p*s>0.17. Free-choice trials cannot differentiate 12-pellet trials vs. 2-pellet trials (i.e., two levers, but a single magazine). There were hardly any magazine entries during these trials. Therefore, magazine entries will not be further discussed in the pSNC task.

Thus, chemogenetic inhibition of the dHC yielded no evidence of an effect on consummatory behavior in the cRD task after a 32-to-2% sucrose downshift, a result consistent with previous reports (see Introduction for references). Following the criterion outlined previously (i.e., 0.33<BF<3; Keysers et al., 2020), Bayes analyses supported a null effect for the Group by Session interactions, although they were inconclusive with a Group effect. However, the same animals showed perseverance of a preference for the option previously associated with the largest reward after a 12-to-2 pellet downshift in the pSNC task based on anticipatory behavior. This result is consistent with similar effects of HC lesions in experiments involving instrumental behavior (see Introduction for references).

## Experiment 3: HC levels of tumor necrosis factor-α (TNF-α) and open field (OF) behavior

The systemic and intracerebroventricular administration of lipopolysaccharide (LPS), an endotoxin derived from the outer membrane of Gram-negative bacteria, induces peripheral inflammation (Page et al., 2022). Several molecular products of inflammation cross the blood-brain barrier and affect neural function to induce sickness behavior (i.e., lack of activity in animals suffering an infectious disease; Hart & Hart, 2019) and cognitive impairments traced to hippocampal dysfunction, including neurogenesis (Kranjac et al., 2012; Valero et al., 2014; Zhao et al., 2019). Some products of inflammation include TNF-α, interleukin-1β, prostaglandin E2, cyclooxygenase-2, brain-derived neurotrophic factor (BDNF), zinc finger (Zif)-268 mRNA expression, and inducible nitric oxide synthase. While these pro-inflammatory molecules propagate throughout the brain, their proliferation has been shown to be greater in the dHC compared to other areas (Kranjac et al., 2012). The effects of LPS-induced inflammation therefore has shown to acutely disrupt hippocampal CA3 and CA1 neural circuit activity (Czerniawski & Guzowski, 2014) and blunt production of hippocampal BDNF, a protein associated with hippocampal neurogenesis (Kranjac et al., 2012, 2013). Cognitive impairments from LPS have also been shown to be HC dependent. These include deficits in water-maze performance, passive and active avoidance learning, and contextual fear conditioning, all hippocampal-dependent tasks, but the same manipulation does not affect hippocampal-independent tasks, such as fear conditioning to a discrete signal (e.g., Pugh et al., 1998; Barrientos et al., 2002). Thus, the use of LPS allows for an additional means of manipulating hippocampal function.

In preparation to study the effects of LPS administration on cRD (Experiment 4) and pSNC (Experiment 5), Experiment 3 aimed to determine the appropriate dosage of LPS that both minimizes sickness behaviors and maximizes cytokine production in the HC. Higher doses of LPS have been shown to decrease locomotor activity, reduce body weight, and facilitate conditioned taste aversion all as a response to inducing a state of illness (Bishnoi et al., 2022; Hart & Hart, 2019; Savi et al., 2021). Therefore, a lower dosage was needed in order to minimize these effects in order to more selectively manipulate neuronal function in the absence of sickness-related behavioral effects. Therefore, Experiment 3 assessed the locomotion effects of a single LPS injection in naïve animals divided into two cohorts (Figure 3a). Cohort 1 was used to select the LPS dose based on the motor effects as recorded in a single OF session (Figure 3b, c). The whole HC was then extracted and cytokine binding was assessed (Figure 3d). Dosage was selected to minimize effects on locomotor activity due to sickness behaviors while still causing a detectable increase in cytokine production in the HC. The selected dose was then used with the second cohort and also to assess LPS effects on cRD (Experiments 4) and pSNC (Experiment 5). Cohort 2 was used to distinguish between dHC and ventral HC (vHC) tissue in terms of cytokine binding (Figure 3e). These animals were not subjected to any behavioral testing.

**Fig 3.**
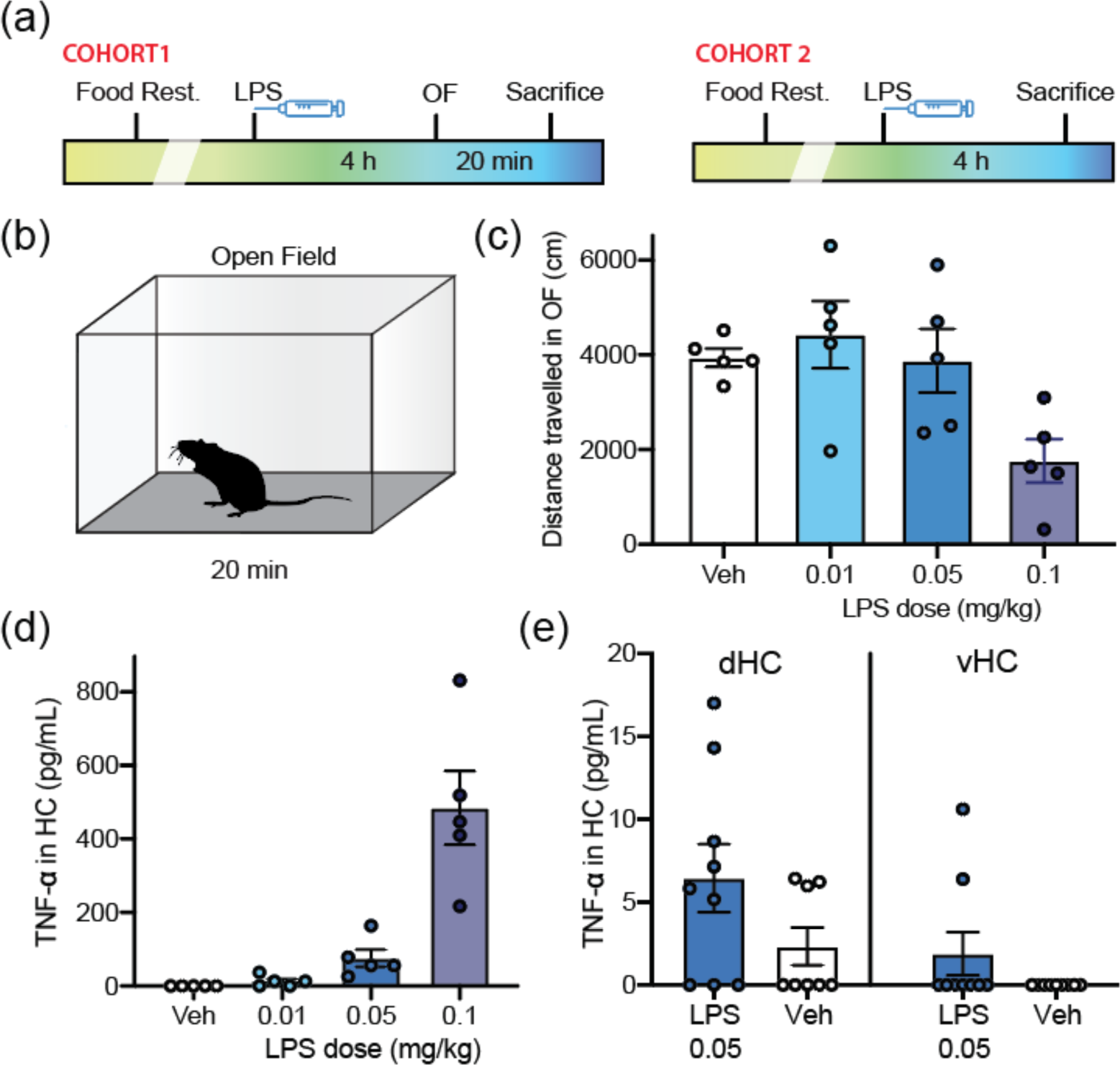
Locomotor and hippocampal inflammatory effects of LPS systemic administration were dose-dependent. (a) Experimental timelines for cohort 1 (*n*=5 per group) and cohort 2 (*n*=9 per group). (b) Illustration of the open-field (OF) apparatus used for animals in cohort 1. (c) Distance traveled in the OF after injections of different doses of LPS or vehicle (cohort 1). (d) Concentration of TNF-α in the HC after injections of different doses of LPS or vehicle (cohort 1). (f) Concentration of TNF-α in the dorsal and ventral HC after injections of LPS (0.05 mg/kg) or vehicle (cohort 2). See text for further details.

## Method

### Subjects

The first cohort involved 20 female Wistar rats with a mean ad libitum weight of 327.8 g (±4.5). The second cohort involved 17 female Wistar rats with a mean ad libitum weight of 250.8 g (±5.7). All these animals were experimentally naïve and approximately 90 days old at the start of the experiment. Although food restriction was not needed for these animals, it was implemented to keep conditions constant with the contrast experiments involving LPS manipulations (Experiments 4-5). Access to food was gradually restricted until animals reached 81-84% of their ad libitum weight. All other conditions were as described in Experiment 2.

### LPS preparation

LPS (*Escherichia coli* serotype 055:B5; Sigma, St. Louis, MO) was diluted in 0.5 mL of sterile, pyrogen-free 0.9% saline (Baxter, Deerfield, IL). The first cohort of animals was injected with Veh (saline, equal volume, 0.2 mL), 0.01, 0.05, and 0.1 mg/kg of LPS. The selected dose, 0.05 mg/kg, was injected to animals in the second cohort. All injections were administered 4 h before tissue extraction.

### OF testing

Animals in cohort 1 were exposed for 20 min to the OF 4 h after receiving an LPS injection (Figure 3a). OF testing was carried out in four units (43×30×43 cm, L×H×W; MED Associates, St. Albans, VT, USA; Figure 3b), illuminated by two 150 W LUMAPRO Clamp-On lights (Grainger, Fort Worth, TX) with 15-W LUMAPRO LED warm white lamp bulbs (Grainger, Fort Worth, TX). A Color Gigabyte Ethernet Camera (Noldus, Leesburg, VA) was placed directly above the open fields and connected to EthoVision XT Version 11 Software (Noldus, Leesburg, VA) to record the activity of animals.

### Cytokine quantification

Animals in cohort 1 were sacrificed immediately after OF testing. The whole HC was extracted and stored in a protein extraction solution containing protease inhibitors (PRO-PREP, Bulldog Bio, Portsmouth, NH), and snap-frozen on dry ice. Samples were stored at -80 °C until processing and analysis. Lysates were centrifuged (16,820 x g) for 20 min and clear lysate was collected.

Animals in cohort 2 were injected with LPS (0.05 mg/kg) or Veh and 4 h later were perfused. The HC was extracted and split evenly into dorsal and ventral sections. Tissue was stored in a protein extraction solution containing protease inhibitors (PRO-PREP, Bulldog Bio, Portsmouth, NH), and snap-frozen on dry ice. Samples were stored at -80 °C until processing and analysis. Lysates were centrifuged (16,820 x g) for 20 min and clear lysate was collected.

Cytokine levels for all samples were measured using TNF-α ELISA kits (BioLegend, San Diego, CA, USA) according to the manufacturer’s instructions.

### Statistics

The dependent variable in the OF test was the total distance traveled during the 20 min measured in centimeters. The dependent variable for the assessment of TNF-α concentration in both cohorts was the value given by the ELISA kit. Many of the samples yielded no TNF-α presence and therefore all the data were analyzed using nonparametric tests, either Kruskal-Wallis for 4 samples or Mann-Whitney for pairwise comparisons of independent samples. A *p*<0.05 was used in all tests.

## Results

The results of the OF test for animals in cohort 1 are presented in Figure 3c. An analysis of all four groups with the Kruskal-Wallis test indicated a significant difference, *χ*(3) =9.17, *p*<0.03. Mann-Whitney pairwise tests indicated that Veh did not differ from animals receiving 0.01 or 0.05 mg/kg, *U*s>5, *p*s>0.17, but activity was significantly higher after Veh than after 0.10 mg/kg of LPS, *U*=0, *p*=0.008. Whole-HC cytokine assessment for these animals is presented in Figure 3d. A Kruskal-Wallis test for the four conditions indicated a significant effect, *χ*(3)=17.10, *p*<0.001. Pairwise Mann-Whitney tests indicated that Veh differed significantly from 0.10 and 0.05 mg/kg, *U*s=0, *p*s<0.006, but not from 0.01 mg/kg of LPS, *U*=5, *p*=0.054. Based on these results, the chosen LPS dose for cohort-2 animals (and for Experiments 4-5) was 0.05 mg/kg.

Cohort 2 assessed whether there was a difference in cytokine presence in the dHC and vHC. The results are presented in Figure 3e. Mann-Whitney pairwise tests were computed for each hippocampal region, but none of the comparisons were significant, *U*s>22, *p*s>0.14. There was also a nonsignificant trend toward higher TNF-α presence in dHC than in vHC, *U*=22.5, *p*=0.081 (Figure 3e).

## Experiment 4: LPS and behavior in the cRD task

The results of Experiment 3 indicate that a single LPS administration (0.05 mg/kg) leads to significant levels of TNF-α in the HC, particularly in the dorsal region. In addition, LPS is known to affect HC function and its behavioral correlates in a set of learning tasks, including contextual fear conditioning (Kranjac et al., 2012), the Morris water maze spatial task (Valero et al., 2014), and passive avoidance (Zhao et al., 2019). The present experiment examined LPS effects on the cRD task, whereas Experiment 5 examined LPS effects in the pSNC task. Based on the results of Experiment 2 with inhibitory DREADDs in the dHC, we expected LPS administration to have no detectable behavioral effects in the cRD task, but to impair behavior in the pSNC task. Manipulating LPS is simpler and less expensive than chemogenetics, therefore allowing for an alternative albeit more indirect approach. However, it is important to note that while LPS has various hippocampal dependent effects both cellularly and behaviorally (see above for references), peripheral injections of LPS leave the possibility for non-hippocampal related effects to emerge. Nevertheless, considering the evidence mentioned previously, it is still hypothesized that LPS will induce similar effects to hippocampal lesions and inactivation. Moreover, given the adapted response to LPS after primary exposure (Grinevich et al., 2001; Erickson & Banks, 2011), the effects of LPS on cRD and pSNC tasks were separated in different experiments and animals only ever given one injection of LPS, rather than different phases and multiple injections as in Experiment 2 with inhibitory DREADDs.

## Method

Twenty female Wistar rats, experimentally naïve and about 90 days of age at the start of the experiment participated in this experiment. Their mean (±SEM) ad libitum weight was 275.8 g (±5.4). The general procedure is described in Figure 4a. The rearing and maintenance conditions, conditioning boxes (Figure 4b), training procedure, and statistical analyses implemented for the cRD task were as described in Experiment 2. LPS preparation and administration were described in Experiment 3.

**Fig 4.**
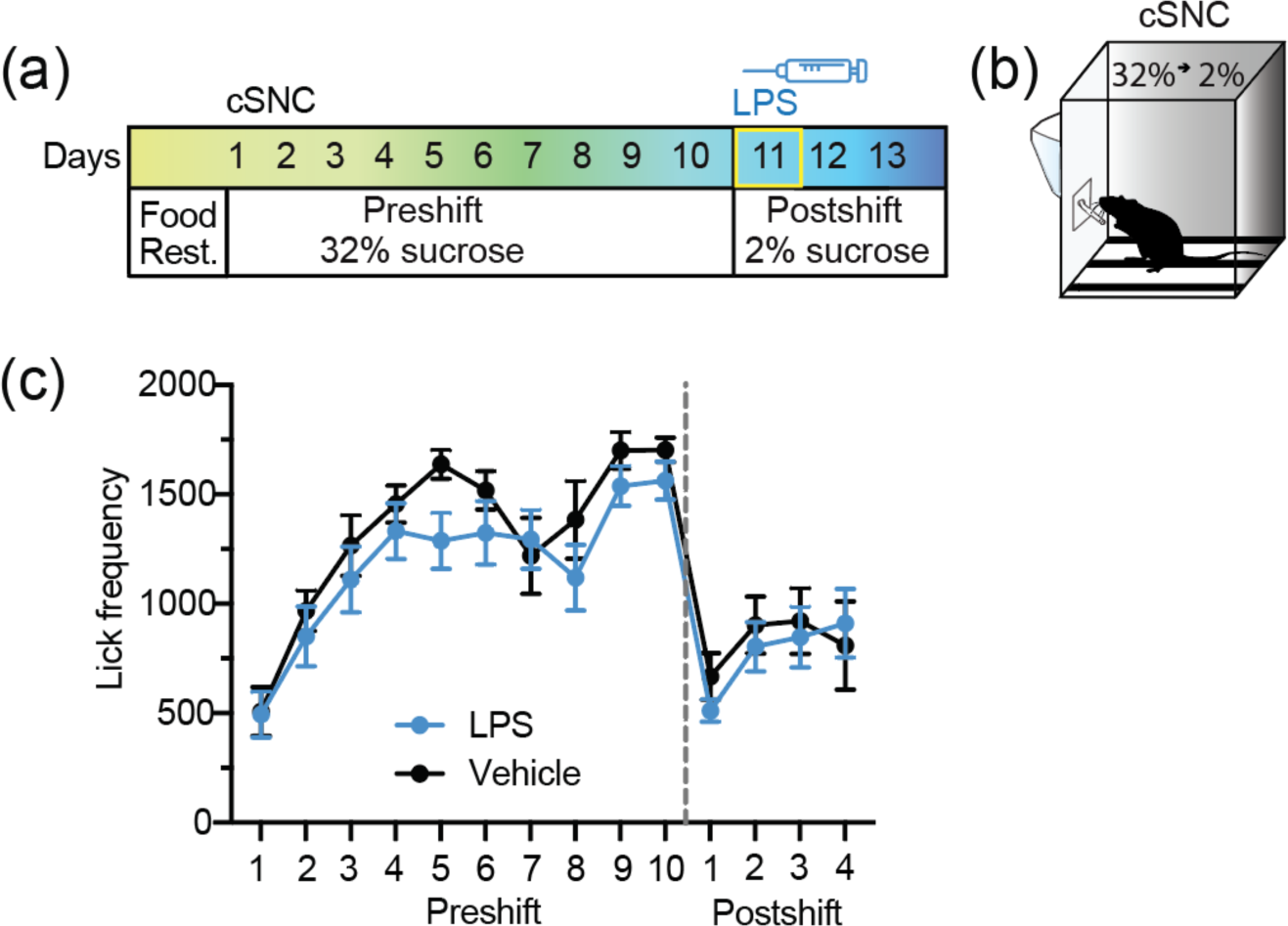
No evidence that peripheral injections of LPS impair adjustment to reward downshifts in a consummatory successive negative contrast task. (a) Experimental timeline. LPS or vehicle were injected intraperitoneally 4h before the beginning of session 11 (first postshift session). (b) Illustration of the behavioral apparatus for the cRD task. (c) Lick frequency during preshift and postshift in the LPS (*n*=10) and vehicle (*n*=10) groups. See text for further details.

## Results

Figure 4c shows the results of this experiment. Animals increased licking for 32% sucrose across preshift sessions 1-10 and exhibited consummatory suppression immediately upon experiencing a downshift to 2% sucrose on postshift sessions 11-13. An analysis of these results failed to reveal significant Group differences (LPS vs. Veh) or a significant Group by Session interaction, either during preshift (Group BF=0.71, Group by Session BF=0.04) or postshift session, *F*s<2.95 (Group BF=0.50, Group by Session BF=0.12). There was a significant increase in licking across preshift sessions, *F*(9, 162)=19.06, *p*<0.001, *η_p_^2^*=0.51, and also a significant recovery of behavior during postshift sessions, *F*(2, 36)=5.86, *p*=0.006, *η_p_^2^*=0.25. An additional analysis indicated a significant suppression of licking from session 10 (last preshift) to session 11 (first postshift), *F*(1, 18)=157.39, *p*<0.001, *η_p_^2^*=0.90, and a marginal Group effect, *F*(1, 18)=4.14, *p*=0.057, *η_p_^2^*=0.19, BF=1.41, but the Group by Session interaction was nonsignificant, *F*<1, BF=0.34. These results are similar to those described in Experiment 2 after the chemogenetic inhibition of dHC neurons.

## Experiment 5: LPS and behavior in the pSNC task

The effects of LPS on the pSNC task were tested separately on preshift and postshift performance. Thus, one group of animals was given LPS before the free-choice test involving the simultaneous presentation of the two levers, one paired with a large reward and the other with a small reward (magnitude test). A second group of animals received the same treatment and magnitude test but was treated with LPS when experiencing a reduction in reward magnitude in one lever, while the other lever remained paired with the small reward (downshift test). This design eliminates potential carry-over effects of LPS from preshift to postshift testing. Based on prior experiments in this series, we expected no detectable effects of LPS on magnitude testing, but a disruption of the normal change in lever preference after reward downshift.

## Method

Sixty-one female Wistar rats, experimentally naïve and about 90 days of age at the start of the experiment participated in this experiment. Their average ad libitum weight was 270.6 g (±3.8). Their rearing and maintenance conditions were those described in Experiment 2. LPS was prepared as described in Experiment 3. Animals were randomly assigned to one of two groups depending on whether LPS was administered before the magnitude free-choice test (Figure 5a) or before the downshift free-choice test (Figure 5d). One set of rats (*n*=30) received LPS or Veh before the magnitude free-choice test. A second set of rats (*n*=31) was injected with LPS or Veh before the downshift free-choice test; these animals were also given a magnitude test, but no LPS or Veh was administered before that session. The rearing and maintenance conditions, conditioning boxes, training procedure, and statistical analyses implemented for the pSNC task (Figure 5b) were as described in Experiment 2.

**Fig 5.**
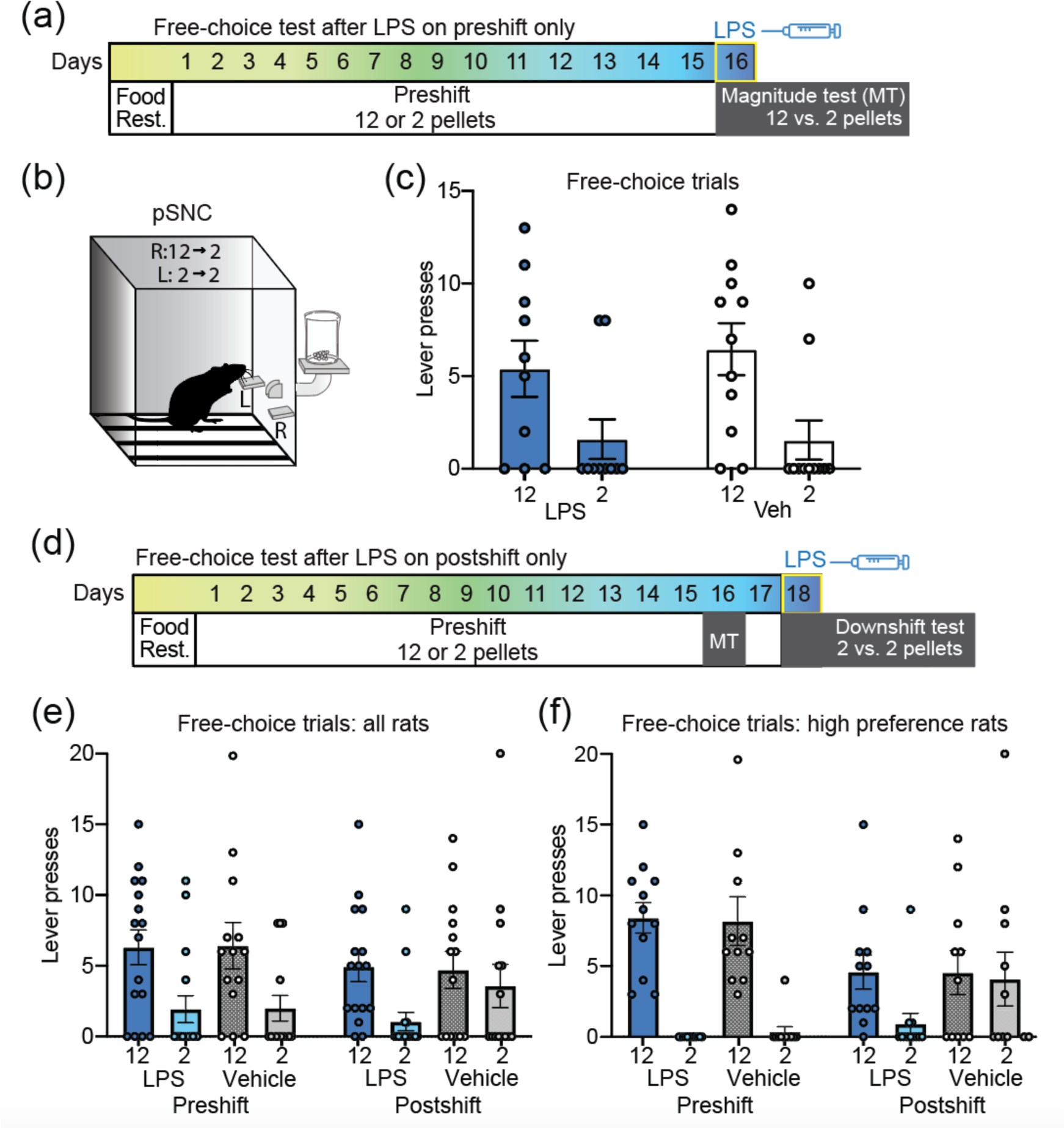
Peripheral injection of LPS impairs adjustment to reward downshifts in a Pavlovian successive negative contrast task. (a) Experimental timeline for group 1. LPS (*n*=10) or vehicle (*n*=11) injections were administered 4 h before the beginning of session 16. (b) Illustration of the behavioral apparatus for the pSNC. (c) Lever presses during the magnitude test (MT) following LPS administration. (d) Experimental timeline for group 2. LPS (*n*=16) or vehicle (*n*=14) injections were administered 4 h before the beginning of session 18. (e) Lever presses during the MT (preshift, no injection), and during the downshift test (postshift, injection) for all rats. (f) Lever presses during the MT (preshift, no injection), and during the downshift test (postshift, injection) for rats that showed a high preference for the large reward during preshift (LPS: *n*=12, vehicle: *n*=11). See text for further details.

## Results

Figure 5c shows the effects of LPS or Veh administration before the nonreinforced free-choice test contrasting levers previously paired with either 12 or 2 pellets. In total, 23/30 rats showed a preference for the 12-pellet lever over the 2-pellet lever, one failed to respond at either lever, and 6 (3 in each group) responded less to the 12-pellet lever than to the 2-pellet lever. Wilcoxon comparisons between the two levers separately for LPS- and Veh-treated animals yielded nonsignificant marginal preference for the 12-pellet lever over the 2-pellet lever in LPS-treated rats, *Z*=-1.792, *p*=0.073, and a significant preference for the 12-pellet lever over the 2-pellet lever in Veh-treated rats, *Z*=-2.078, *p*=0.038. Given the similarity in preference across groups, a Wilcoxon nonparametric test including all the animals was also computed comparing 12- vs. 2-pellet levers. Preference for the lever paired with the largest reward was significant, *Z*=-2.80, *p*=0.005.

A different group of rats was exposed to the same procedure, including the free-choice test for reward magnitude in the absence of the LPS treatment, and then given a reward downshift involving a free-choice trial between the downshifted lever (12-to-2 pellets) and the unshifted lever (always paired with 2 pellets). The results for all the animals are presented in Figure 5e. As with the previous set of rats, preference for the 12-pellet lever over the 2-pellet lever in the preshift free-choice trial was statistically marginal for either group, *Z*s<-1.41, *p*s>0.06. However, because these animals were treated similarly, a second test compared all the animals together. This test yielded a significant preference for the 12-pellet lever over the 2-pellet lever, *Z*=-2.44, *p*=0.015. In the postshift free-choice trial, preference for the lever formerly paired with 12 pellets (downshifted lever) over the lever always paired with 2 pellets (unshifted) was significant for LPS rats, *Z*=-1.86, *p*=0.020, but not for Veh rats, *Z*=-0.723, *p*=0.470.

We also selected for analysis rats that had shown preference for the 12-pellet lever over the 2-pellet lever in the preshift free-choice trial and assessed their preference for the downshifted vs. unshifted lever (Figure 5f). As expected, selected animals in both groups exhibited strong preference for the 12-pellet lever over the 2-pellet lever in preshift, *Z*s>-2.94, *p*s<0.003. Again, for these selected rats, preference for the lever previously paired with 12 pellets (downshifted) was maintained in LPS animals, *Z*=-2.247, *p*=0.025, but that preference was eliminated in Veh animals, *Z*=-0.311, *p*=0.755.

These results showed that the 12-to-2 pellet downshift did not affect a preference for the lever previously paired with 12 pellets over that paired with 2 pellets in LPS-treated rats. However, Veh-treated animals exhibited the usual disruption of the preference after reward downshift. These results in the pSNC task are like those obtained also with anticipatory responses after complete HC lesions (Experiment 1) and inhibitory DREADD infusion in the dHC (Experiment 2).

## Experiment 6: c-Fos expression in dHC and vHC after cSNC

There has been little evidence that the HC is involved in reward downshift in consummatory tasks. The present Experiments 2 and 4 failed to observe behavioral consequences of dHC disruption either after inhibitory DREADDs or LPS treatment, respectively. However, Pecoraro and Dallman (2005) examined the expression of c-Fos, a protein that reflects neuronal activation, and reported increased expression only during the first downshift session and predominantly in ventral areas of the hippocampus (vHC). Some results suggest that there might be a functional differentiation between the dHC, mostly centered on cognitive functions, and the vHC, which appears to be biased toward emotional processing (Fanselow & Dong, 2010; but see Lee et al., 2019). Therefore, Experiment 6 looked for evidence of hippocampal involvement in reward downshift in the DG and CA fields (CA1, CA2-CA3) in both dHC and vHC with a training procedure similar to that used in Experiments 2 and 4, except that two unshifted controls were added: one always exposed to 2% sucrose and the other always exposed to 32% sucrose. The inclusion of unshifted controls justifies describing this as a cSNC effect.

## Method

### Subjects

The subjects were 20 male Wistar rats, experimentally naïve, and 90-day old at the start of the experiment, purchased from Envigo (Barcelona, Spain). Their mean (±SEM) ad libitum weight was 393.9 g (±8.6). Rats were housed individually in polycarbonate cages with water continuously available, in a room with constant temperature (18-22 °C) and humidity (50-60%). Lights were turned on between 08:00 and 20:00 h. Animals were maintained within 82-85% of their ad libitum weight by an appropriate amount of standard rat chow provided at least 20 min after each session. The experiment followed the European Union directive guidelines for the use of animals in research (2010/63/EU) and Spanish Law (6/2013; R.D.53/2013).

### Apparatus

Consummatory training involved six boxes, each measuring 30×15×30 cm (L×W×H), with clear Plexiglas walls, floor, and ceiling. The floor was covered with a layer of sawdust. A sipper attached to a graduated cylinder was inserted through the front wall. The 32% (or 2%) sucrose solution was prepared weight/weight by mixing 32 g (or 2 g) of sucrose for every 68 g (or 98 g) of distilled water. Sucrose was dissolved with a magnetic mixer (Nahita, 680-9, Beriain, Spain). Session length was measured with a manual stopwatch (Extech, 365510, Madrid, Spain).

### Behavioral procedure

Subjects were matched by weight and randomly assigned to Groups 32-2, 2-2, or 32-32 (Figure 6a-b). A 5-min habituation session in the consummatory box without fluids preceded training. Animals had free access to 32% (groups 32-2 and 32-32) or 2% (group 2-2) sucrose on preshift sessions 1-10. One postshift session was administered on day 11, exactly as during preshift sessions for groups 32-32 and 2-2, whereas group 32-2 was downshifted from 32% to 2% sucrose. Each session lasted 5 min from the first contact with the sipper tube. Animals were run in squads of six or eight (all from the same group). On session 11, squads receiving access to 2% sucrose were run first, then those receiving 32-to-2% sucrose, and finally animals exposed to 32-to-32% sucrose. The amount of sucrose solution consumed was transformed to mL/kg based on the animal’s weight recorded on the same day (Figure 6c). Amount consumed, licking frequency, and time in contact with the sipper tube have been previously used as dependent measure in the cSNC task (e.g., Papini, Mustaca, & Bitterman, 1988; Riley & Dunlap, 1979). These dependent measures are also closely related (e.g., Mustaca, Freidin, & Papini, 2002; Spector & Smith, 1984).

**Fig. 6.**
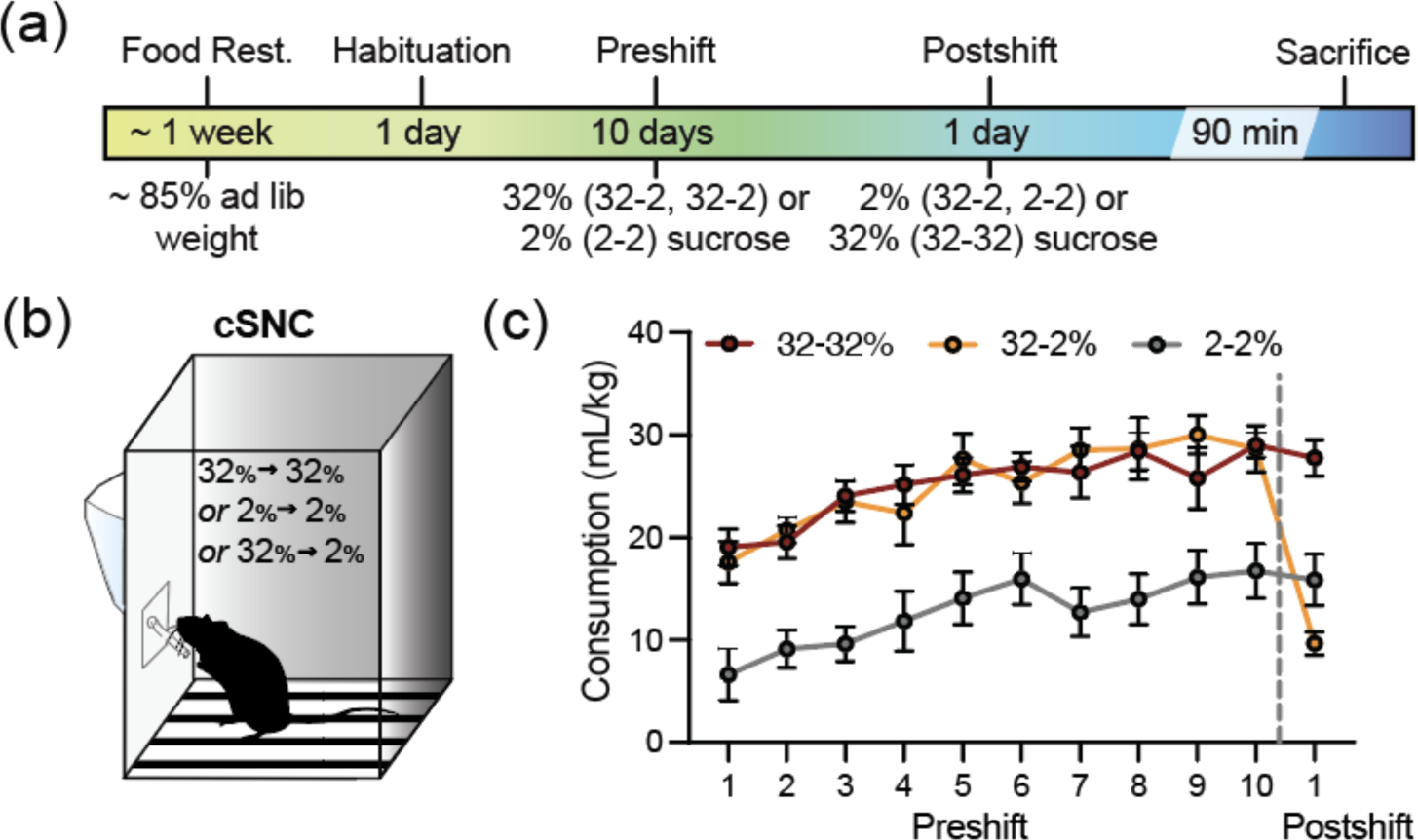
Reward downshift in a consummatory task caused a negative contrast effect. (a) Experimental timeline. (b) Illustration of the behavioral apparatus for the cSNC. (c) Consumption of sucrose (mL/kg) during preshift and postshift in the 32-32 (*n*=6), 2-2 (*n*=6), and 32-2 (*n*=8) groups.

### Histology

Animals were perfused 90 min after session 11 in the cSNC task for maximum c-Fos expression (Morgan & Curran, 1991). Animals received an ip injection of 80 mg/kg of sodium pentobarbital (ABBOT, Madrid, Spain). Once the absence of the foot and tail reflexes was verified, animals were placed in a supine position for the collection of lavage and fixation fluids using a perfusion pump (ISMATEC, MCP/BVP, Wertheim, Germany). After stretching and holding the animal’s limbs, the skin was cut at the end of the sternum, separating the skin and cutting the ribs parallel to the lungs to expose the heart, and a needle was inserted to the left ventricle. The needle was connected to a tube that delivered PBS (0.01 M). An incision into the right atrium allowed for the escape of lavage fluids. Then, saline solution and 10% formalin were infused until the corresponding muscle contraction was observed, indicating the tissue fixation. Finally, the brain was extracted and kept in a 10% formalin solution for 3 h at room temperature and subsequently moved to a 30% sucrose and 0.02% azide solution, and maintained at 4 °C for at least 3 days.

Subsequently, the brains were rinsed in PBS (0.01 M) and then laminated in 30-µm-thick coronal sections with a cryostat (Leica, CM1950, Barcelona, Spain). Every seven sections, one was selected for analysis (Pecoraro & Dallman, 2005). Sections were placed in serial order and maintained free floating in PBS (0.01 M). Brain slices at three A/P coordinates were assessed in each brain: -3.12 (dHC), -4.68 (dHC, vHC), and -5.76 (vHC). For the immunohistochemical analysis of c-Fos, tissue slices were embedded in a 0.3% hydrogen peroxide solution in PBS (0.01 M, pH 7.4) for 30 min at room temperature and in gentle agitation, followed by three washings with PBS (0.01 M) of 5 min each at room temperature and moderate agitation. Next, sections were incubated in 2% bovine serum albumin (BSA) in 0.1% Triton X-100 solution diluted in PBS (0.01 M) for 4 h at room temperature and with gentle shaking. Sections were then incubated overnight at 4 °C temperature with a rabbit polyclonal anti-c-Fos IgG (1:2000; Abcam, Cambridge, United Kingdom), diluted in a 0.2% Triton X-100 solution in PBS (0.01 M) under gentle agitation. Sections were then washed three times, 5 min each time, in PBS (0.01 M), and incubated with a biotinylated goat anti-rabbit IgG(H+L) secondary antibody (1:200; Thermo Fisher Scientific, Massachusetts, USA) diluted in PBS (0.01 M) for 1 h at room temperature with gentle shaking. Sections were washed three times, 5 min each, in PBS (0.01 M) and then incubated with an avidin–biotin–peroxidase complex (ABC; Vector Laboratories, Newark, CA) for 1.5 h at room temperature with gentle shaking. Sections were then washed three times, 5 min each, in an acetate buffer (0.1 M, pH 6.0) at room temperature with moderate shaking. A black chromagen was produced by embedding the sections in a solution containing 3,3-diaminobenzidine tetrahydrochloride (Sigma, Madrid, Spain), 0.4% NH4Cl, 20% b-Dglucose, and 1% nickel ammonium sulfate. All reactions were carried out for 6 min before being stopped by rinsing of the sections three times, 5 min each, in acetate buffer (0.1 M) at room temperature and moderate shaking. After this procedure, the sections were placed on gelatinized slides and allowed to dry for at least 24 h. Sections were subsequently dehydrated and rinsed with alcohol of increasing concentrations (50%, 70%, 96%, 100%) and xylol (I, II) for 1 min in each solution. Sections were mounted and covered with DPX, and dried for at least 24 h. Sections were visualized at the Optical Microscopy Unit, University of Jaén, with a transmitted light laser (488 nm) of fluorescence inverted microscope (DM IRB, Leica Biosystems, Wetzlar, Germany), and an inverted confocal microscope (TCS SP5II, Leica Biosystems, Wetzlar, Germany). Photomicrographs of dorsal and ventral hippocampal regions (DG, CA1, and CA2-CA3) were taken at 10 x magnification using either a C-MEX-18 Pro camera (Euromex Microscopen, Arnhem, The Netherlands) or the above-mentioned confocal microscope.

Histological counting of c-Fos positive cells in the brain regions of interest was obtained using the ImageJ 1.54 b Software (National Institute of Mental Health, Maryland, USA). c-Fos-positive cells were automatically identified by the software by thresholding objects with 0.98-1.00 circularity value, matching c-Fos positive nuclei. To minimize background noise and equalize all the microphotographs they were previously converted into 8-bit-type images and lightened (50.0 pixels).

### Statistics

In accordance with previous studies (e.g., Agüera et al., 2023; Donaire et al., 2022; Ruiz-Salas et al., 2022), the amount of sucrose consumed was normalized based on body weight (mL/kg) for a more meaningful comparison across individuals and across groups. The density of c-Fos positive cells relative to the area in which they were recorded (cells/mm^2^) was also calculated. These variables were assessed with conventional analysis of variance, with a *p*<0.05 level for significance; *η_p_^2^* assessed effect size, and pairwise Bonferroni tests identified the source of significant interactions.

## Results

Figure 7a shows representative photos of the DG for each of the three reward conditions, 32-2, 2-2, and 32-32. The number of hippocampal c-Fos positive cells was generally low. Figure 6c shows the behavioral results for the cSNC task in terms of sucrose consumption (mL/kg) on sessions 10 (last preshift) and 11 (first postshift). Three results were of interest. First, unshifted groups (2-2 and 32-32) did not change their consumption from preshift to postshift. Second, rats consume more 32% sucrose than 2% sucrose. Third, reward downshift caused a drastic suppression of performance on session 11 relative to session 10 in Group 32-2. A Group (2-2, 32-32, 32-2) by Session (1-11) analysis indicated significant effects for all factors and their interaction *F*s>6.63, *p*s<0.01, *η_p_^2^*s>0.43. Bonferroni pairwise tests indicated higher sucrose intake in groups receiving 32% sucrose (32-32 and 32-2) compared with Group 2-2 in preshift sessions (ps<0.05), and a significant reduction in sucrose consumption from preshift sessions 2-10 to postshift session 11 in Group 32-2, *ps*<0.002, but no evidence of change across sessions in Groups 2-2 and 32-32, *p*s>0.06.

**Fig. 7.**
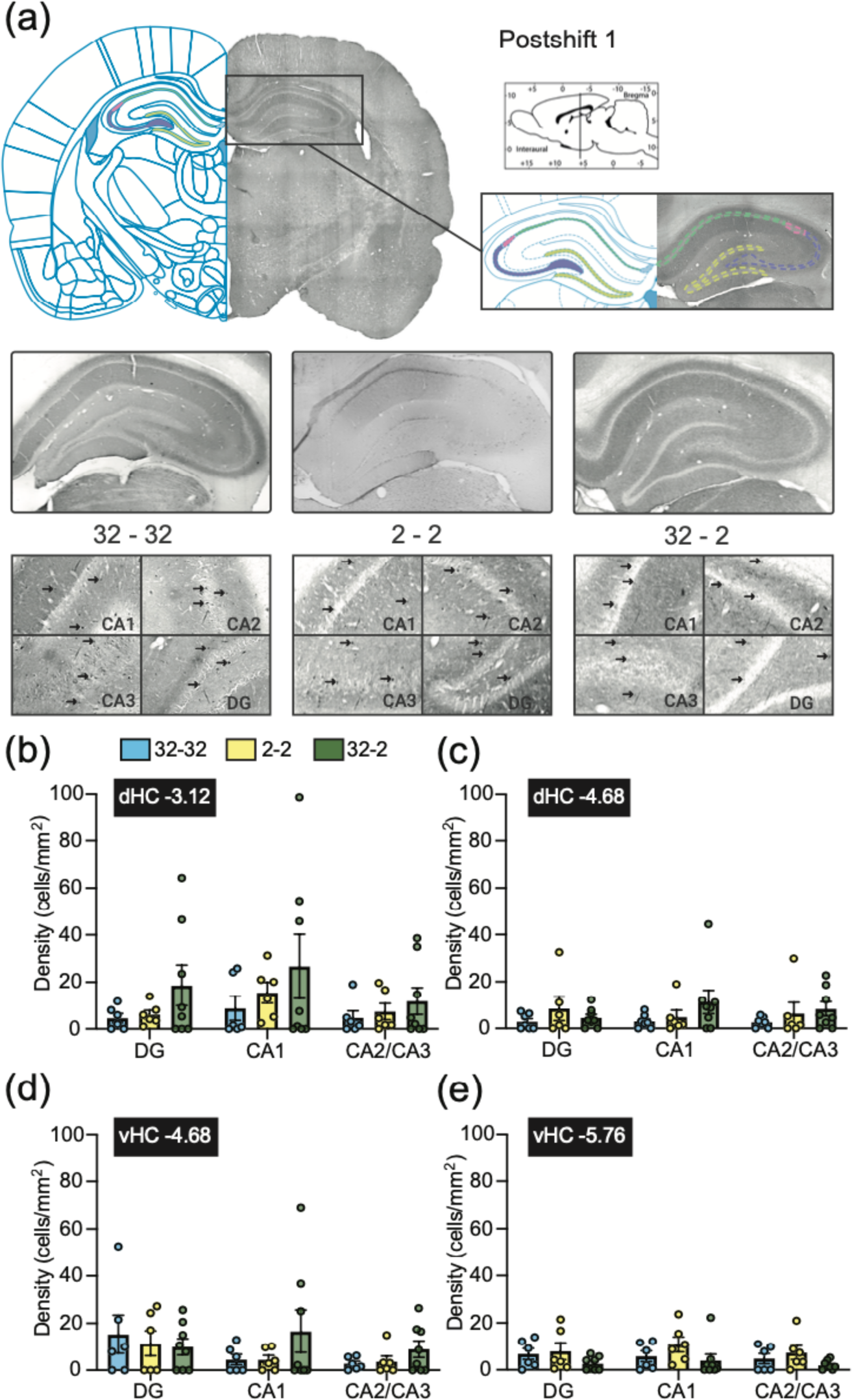
No evidence that reward downshift in a consummatory task caused an increase in c-Fos expression in the dorsal or ventral hippocampus (dHC, vHC). (a) Representative examples of brain section photographs for each group (A/P coordinate -3.12). Arrows indicate examples of c-Fos positive cells. (e-h) c-Fos cell density in the dorsal (e, f) and ventral (g, h) DG, CA1, and CA2/3 regions of the HC for each group (A/P coordinates on the top-left corner of each figure). See text for further details.

Figures 7b-e show the density of c-Fos positive cells in each of the slices selected for analysis along the A/P axis. Independent Group (32-32, 2-2, 32-2) by Area (DG, CA1, CA2-CA3) analyses were computed for each A/P coordinate. In none of the analyses was the interaction or the Group effect significant, *F*s<1.84, *p*s>0.14, *η_p_^2^*s>0.18. The Area effect was significant only for the A/P coordinate -3.12, dHC, *F*(2, 34)=4.45, *p*<0.02, *η_p_^2^*=0.21, but Bonferroni comparisons indicated that none of the pairwise tests was significant, *p*s>0.10. Because the cell counts in these HC regions were relatively low, we looked at the habenula (both lateral and medial) as a positive control and found higher densities in the same animals. Mean cell densities were 35.7 for Group 32-32, 28.5 for Group 2-2, and 79.7 for Group 32-2.

These results are consistent with the lack of behavioral differences in cSNC observed after hippocampal lesions (Flaherty et al., 1989, 1998; Kramarcy et al., 1973), inhibitory DREADDs (Experiment 2), and LPS administration (Experiment 4). These c-Fos results provide no clear support for the possibility that chemogenetic inhibition of neural activity in the vHC during cSNC would disrupt behavior in the cSNC task. Therefore, the tentative conclusion is that there is no evidence that the cSNC task engages hippocampal circuits.

## General discussion

The capacity of organisms to adjust their behavior based on past experience with shifting reward conditions is fundamental to their fitness. Understanding the neural mechanisms that drive these behavioral changes is essential for an understanding of interplay between memory systems and adaptive behavior. The HC is central to these mechanisms, playing a pivotal role in shaping behaviors based on experience. Yet, conflicting findings from studies on reward downshift and HC function have left unresolved questions. Specifically, how crucial is the HC in guiding behavioral modifications in response to changes in reward? Does HC function depend on the situation or on the specific actions an animal undertakes to obtain the reward? We examined these questions in a comprehensive set of experiments aiming at elucidating the contribution of the HC to behavioral adjustments following reward downshifts.

Two experiments revealed no effects of neural manipulations on the cRD task after chemogenetic inhibition of the dHC (Experiment 2) and LPS manipulations that influenced dHC cells (Experiment 4). These results aligned with the absence of evidence for increased dHC and vHC activation, measured by c-Fos expression, following a reward downshift in the cSNC task (Experiment 6). However, reward devaluation 8-maze task and pSNC tasks, where animals freely choose between downshifted or unshifted reward options, HC dysfunction influenced behavior, whether after complete excitotoxic lesions (Experiment 1), chemogenetic inhibition of the dHC (Experiment 2), or LPS treatment impacting the dHC (Experiment 5). The similarity of the results between these experiments is striking. This is especially relevant with Experiments 4 and 5 given the systemic nature of LPS-induced inflammation. These congruent results give further credibility for the use of LPS as an additional manipulation of the HC function.

The distinction between consummatory and anticipatory tasks (whether instrumental or Pavlovian) regarding the involvement of the HC is noteworthy. The present results provided no evidence that consummatory behavior after reward downshift relies on the HC. However, evidence of enhanced c-Fos expression in the CA3 region, entorhinal cortex, postsubiculum, and ventral subiculum in the cSNC task was reported by Pecoraro and Dallman (2005). This study reported elevated c-Fos expression in these HC areas after the first downshift session. The only region that overlaps with the present c-Fos data is the CA3 field and the present findings failed to confirm these previous results. There were two procedural differences in training that might be responsible for this discrepancy. Pecoraro and Dallman (2005) used a 32-to-4% sucrose downshift and 12 preshift sessions, rather than 32-to-2% sucrose downshift and 10 preshift sessions as used in Experiment 6. None of these differences appears a priori to justify differential CA3 c-Fos expression. However, they assessed CA3 separately from CA2, whereas we combined these two areas into a single score because of the relatively low level of c-Fos expression (see Figure 7b-e). Despite this inconsistency in c-Fos data in the CA3, and given the support for a null Group by Session interaction effect for postshift performance from Bayes analyses in both Experiments 2 and 4, we suggest that reward downshifts effects are independent of HC function. This conclusion is also consistent with earlier reports of no evidence that HC lesions affect the cSNC effect (Flaherty et al., 1989, 1998; Kramarcy et al., 1973).

Unlike consummatory tasks, behavioral adjustments in tasks requiring anticipatory responses to changes in reward value necessitate the HC. Compared to cRD and cSNC tasks, the anticipatory tasks used in this series (Experiments 1, 2, and 5) present several differences. First, the reward locations were spatially differentiated, requiring the animal to either move right or left in the 8-maze or orient toward right or left levers in the pSNC task. The cSNC task requires animals to locate the source of reward, but reward location is consistent across the experiment, a fact that would make a small demand for spatial orientation. The observed sensitivity of anticipatory tasks to HC manipulations highlights the HC’s pivotal role in behaviors linked to spatial navigation and reward location recognition. Such a role is consistent with prior studies demonstrating that HC neurons encode spatial information and update these representations based on reward cues (Dupret et al., 2010; O’Keefe, 1976; Poucet & Hok, 2017).

Second, if behavioral changes mirror the modification of appetitive memories, then the observed resistance to change in the anticipatory behavior of animals with a non-functional HC could indicate a deficit in memory updating. This aligns with multiple studies that emphasize hippocampal retrieval-induced plasticity as central to the reconsolidation process, essential for adjusting existing memories (Lee, 2009, 2010; Sara, 2000; Winters et al., 2009). As in previous similar studies (e.g., Franchina & Brown, 1971), our results showed no evidence that HC integrity is necessary for the initial acquisition of differential responding as a function of reward magnitude, but HC integrity is required when the original memory needs updating after a reward downshift.

Third, the anticipatory tasks implemented in this series also require animals to confront two reward options simultaneously in free-choice trials. There are indications that, both in human and non-human animals, the HC is involved when decisions are based on the value of the options (Amemiya & Redish, 2016; Johnson et al., 2007; Palombo et al., 2015; Papale et al., 2016). Bakkour et al. (2019) reported that participants choosing between images (signals) of food items that varied in value (e.g., pictures of two different chocolate bars) exhibited longer reaction times when the items were closer in value than when one item was far more preferred than the other. Importantly, reaction times were positively correlated with BOLD (blood-oxygen-level-dependent) activity in the HC using functional magnetic resonance imaging (fMRI) in healthy participants. Furthermore, amnesic patients with medial temporal lobe damage exhibited longer reaction times when making decisions about signals for rewards of different value. While it is possible that the effects reported in the present anticipatory tasks are related to choosing between options differing in the reward values they signaled, this mechanism should have led to measurable effects in preshift performance. Such preshift effects of HC manipulations were not found in the present tasks (see Figures 1e, 2e, and 5e-f). In addition, we observed that hippocampal lesions resulted in decreased VTE behavior (Figure 1f), a phenomenon that has been associated with the deliberative processes in decision-making (Johnson et al., 2007), suggesting a potential impairment in the ability to simulate or deliberate over future paths. These results are in alignment with previous research demonstrating a critical role of the HC in decision-making and the modulation of anticipatory behaviors (Amemiya & Redish, 2016; Johnson et al., 2007).

Fourth, there is compelling evidence indicating that, in tasks involving reward downshifts, consummatory and anticipatory behaviors have distinct characteristics, regardless of HC function. Behaviorally, a study replicating the pSNC effect with consummatory behavior showed a preference for the option offering the highest sucrose concentration, mirroring findings using the autoshaping procedure (Conrad & Papini, 2018). However, unlike in the pSNC task, this preference remained unaltered following a reduction in sucrose concentration (Guarino et al., 2020). Moreover, in runway training experiments with a sucrose downshift presented in the goal box, evidence was found for cSNC, indicated by a decrease in sucrose consumption within the goal box. However, there was no evidence of iSNC in the runway leading up to the goal box, as the latency to reach the goal box remained unchanged (e.g., Sastre et al., 2005). From a neurobiological perspective, lesions of the nucleus accumbens abolish the iSNC without affecting the cSNC (Leszczuk & Flaherty, 2000), whereas lesions of the gustatory thalamus produce the opposite impairment (Sastre & Reilly, 2006). Our distinct results using HC manipulations emphasize the importance of the differentiation between consummatory and anticipatory behaviors following reward downshifts, and they further suggest a more context or response-specific engagement of the HC than previously recognized (Dupret et al., 2010; O’Keefe, 1976; Poucet & Hok, 2017).

Fifth, these tasks do possess some distinct procedural elements. Specifically, the anticipatory tasks in this series utilized solid rewards and implemented multiple trials per session. By contrast, the cSNC task involved liquid rewards and a single trial per session. However, these differences do not appear to be the defining factors that set the tasks apart when considering them independently of the influence of the HC. For example, cSNC has been reported to lead to similar behavioral changes whether using sucrose solutions or solid food rewards (Pellegrini & Mustaca, 2000), and the iSNC effect was reported many times with training involving a single trial per session (one session per day), including the initial reports using a complex maze (Elliott, 1928) and a runway with rats (Crespi, 1942).

This series of experiments provides insight into the function of the HC in adjusting behavior to reward downshifts, underscoring its functional relevance in specific reward downshift tasks. The dHC is engaged when reward choice requires spatial identification of response locations, when a change in reward value must be followed by the update of the relevant memory, when there is a decision to be made between rewards differing in value, and/or when an animal must make a response in anticipation of a change in reward value.

Ideally, HC manipulations could be tested in consummatory and anticipatory tasks within the same experiment. However, integrating these tasks poses significant challenges due to the complex interplay of cognitive processes involved. An alternative approach would be to identify dimensions that differ between these tasks and test the effects of HC manipulations in one type of task. In addition to the response type, consummatory and anticipatory tasks also differ in the degree of spatial dependency, the response complexity, and the degree of response flexibility, higher for anticipatory than consummatory tasks in all three cases. For example, response complexity could be increased in the consummatory situation by introducing a free-choice component, thus offering access to bottles containing different sucrose concentrations (e.g., Guarino et al., 2020). Moreover, the degree of response flexibility could be assessed by reversing the position of the bottles in a second phase. If these dimensions are more relevant than the type of response involved, then HC lesions or chemogenetic inhibition should disrupt free-choice consummatory performance and retard or prevent the reversal of preference relative to controls. This remains to be tested.

By revealing these nuanced roles of the dHC in guiding goal-oriented actions, we advance our understanding of how animals adjust to changes in reward conditions. More broadly, these data also emphasize the importance of considering the specific properties of paradigms and contexts when investigating neural circuits of behavioral phenomena.

## Acknowledgments

Partial funding for this research was obtained from the following sources: Mount Holyoke College (MHC) Program in Neuroscience and Behavior, as well as the MHC Faculty Research Support Grant (to MS); the Curtis Smith Award (to MH); SERC Grant 210321 (to CH); NIDA Drug Supply Program (to MRP); TCU IS Grant 66054 (to MRP); SERC Grant (to MRP); Spanish Ministerio de Ciencia e Innovación Grant PID2021-123338NB-100 (to CT); and the “Margarita Salas Fellowship,” Grant UJAR07MS, Spanish Ministry of Universities (to PO). We thank L. Anderson, D. Chohan, S. Guarino, R. Rodríguez, A. Sánchez, D. Scepanovic, F. Vignolo, E. Vogiatzoglou, and D. Zafra for assistance in several stages of the research presented here. P. Ogallar’s current affiliation is Psychological and Brain Sciences Department, University of Iowa, Iowa City, IA 52242, USA. Corresponding author: M. Sabariego.

## Conflict of interest statement

Authors declare no conflicts of interest

## Highlights

- Reward downshifts were implemented in consummatory, instrumental, and Pavlovian tasks
- HC lesions, chemogenetic inhibition, and LPS treatments interfered with anticipatory behavior
- No evidence that interfering with HC function affected consummatory behavior
- No evidence of differential c-Fos expression in consummatory successive negative contrast
- HC interference selectively affected tasks requiring reward anticipation and choice

## CRediT authorship contribution statement

**C. Hagen:** conceptualization, investigation, data curation, formal analysis, funding acquisition, writing: review and editing. **M. Hoxha:** investigation, data curation, funding acquisition, writing: review and editing. **S. Chitale:** investigation, writing: review and editing. **A. O. White:** investigation, writing: review and editing. **P. Ogallar:** investigation, data curation, writing: review and editing. **Expósito, A.N.:** investigation, data curation, visualization, writing: review and editing. **A. D. R. Agüera:** investigation, data curation, writing: review and editing. **C. Torres:** conceptualization, data curation, formal analysis, funding acquisition, methodology, project administration, supervision, visualization, writing: original manuscript, review, and editing. **M. R. Papini:** conceptualization, data curation, formal analysis, funding acquisition, methodology, project administration, supervision, visualization, writing: original manuscript, review, and editing. **M. Sabariego:** conceptualization, data curation, formal analysis, funding acquisition, methodology, software, project administration, supervision, visualization, writing: original manuscript, review, and editing.

## Significance statement

Hagen et al. elucidate the specific role of the hippocampus (HC) in adjusting behavior when rewards are devalued—a key adaptive challenge. By conducting novel experiments with rats, the research reveals the HC’s crucial, yet nuanced role in shaping anticipatory behaviors—not immediate consumption—in response to reward reduction. These findings untangle previous contradictions about the HC’s specific functions, highlighting its targeted role in preparing for future rewards rather than immediate consumption. The results offer clear implications for understanding adaptive behavior and inform potential treatments for reward-related disorders, marking a significant advance in neuroscience and psychology.

## Supplementary Figures

**Figure S1.**
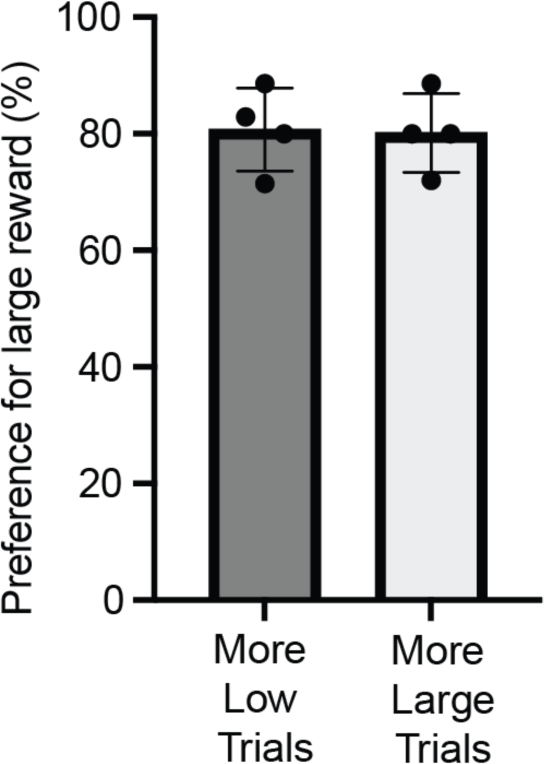
Preference for the large reward remained unaffected by variations in force trail numbers. During preshift testing, rats consistently chose the large reward location over the low reward option in free-choice trials, a pattern that was observed across all preshift testing sessions. Varying distribution of force trials, where rats were directed to a specific reward location, did not alter this preference.

**Figure S2.**
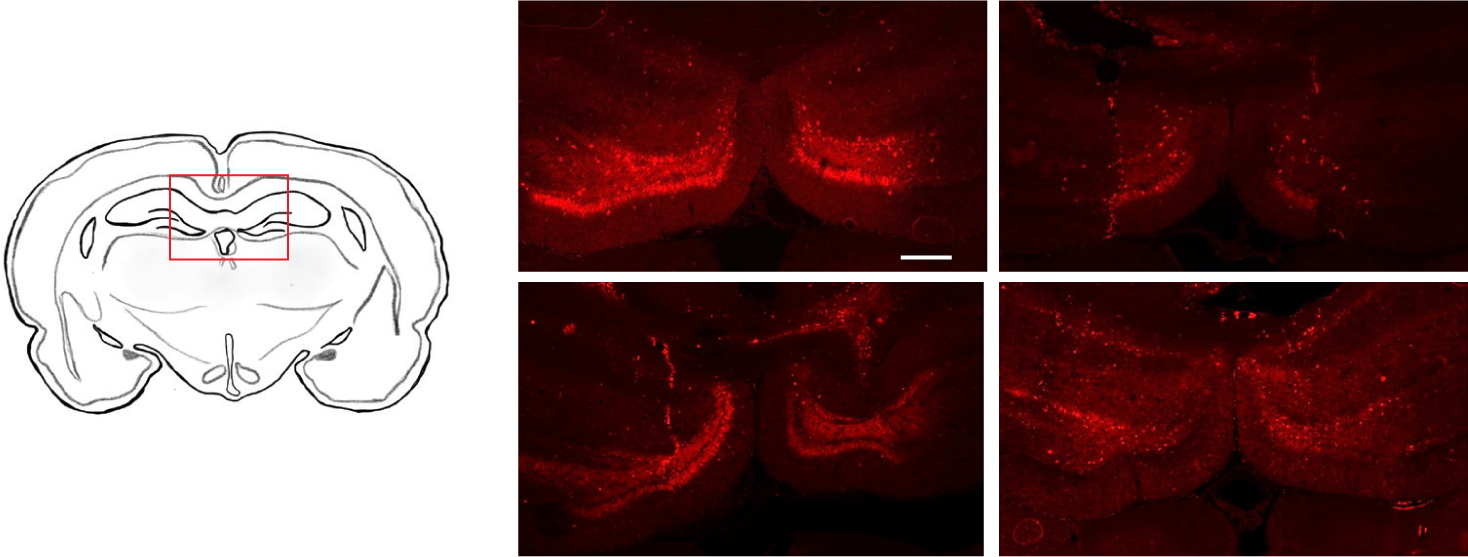
DREADD Expression in dorsal hippocampus across different subjects. Representative coronal brain sections from four different animals illustrating the extent of AAV2-hSyn-hM4D(Gi)-mCherry viral expression. The expression is predominantly localized within the dentate gyrus (DG) of the dorsal hippocampus (dHC), as evidenced by the mCherry fluorescence. Scale bar: 500 μm

## Notes

### Competing Interest Statement

The authors have declared no competing interest.

